# Rhabdomyosarcoma fusion oncoprotein initially pioneers a neural signature in vivo

**DOI:** 10.1101/2024.07.12.603270

**Authors:** Jack Kucinski, Alexi Tallan, Cenny Taslim, Meng Wang, Matthew V. Cannon, Katherine M. Silvius, Benjamin Z. Stanton, Genevieve C. Kendall

## Abstract

Fusion-positive rhabdomyosarcoma is an aggressive pediatric cancer molecularly characterized by arrested myogenesis. The defining genetic driver, PAX3::FOXO1, functions as a chimeric gain-of-function transcription factor. An incomplete understanding of PAX3::FOXO1’s in vivo epigenetic mechanisms has hindered therapeutic development. Here, we establish a PAX3::FOXO1 zebrafish injection model and semi-automated ChIP-seq normalization strategy to evaluate how PAX3::FOXO1 initially interfaces with chromatin in a developmental context. We investigated PAX3::FOXO1’s recognition of chromatin and subsequent transcriptional consequences. We find that PAX3::FOXO1 interacts with inaccessible chromatin through partial/homeobox motif recognition consistent with pioneering activity. However, PAX3::FOXO1-genome binding through a composite paired-box/homeobox motif alters chromatin accessibility and redistributes H3K27ac to activate neural transcriptional programs. We uncover neural signatures that are highly representative of clinical rhabdomyosarcoma gene expression programs that are enriched following chemotherapy. Overall, we identify partial/homeobox motif recognition as a new mode for PAX3::FOXO1 pioneer function and identify neural signatures as a potentially critical PAX3::FOXO1 tumor initiation event.

## Main

Rhabdomyosarcoma is a pediatric cancer defined by the presence or absence of gene fusions, with fusions generally predictive of a worse clinical prognosis^1–4^. In the fusion-positive rhabdomyosarcoma (FP-RMS) subtype, the most common fusion oncogenes are *PAX3::FOXO1* and *PAX7::FOXO1*, which juxtapose the paired-box and homeobox DNA binding domains of PAX3/7 to the transactivation domain of FOXO1^5–8^. The PAX3/7::FOXO1 oncoproteins function as aberrant transcription factors with gain-of-function activities^9–12^. Despite the discoveries of PAX3/7::FOXO1 over three decades ago, there are no targeted therapies for the disease^13^. FP-RMS is less responsive to the currently used treatments of chemotherapy, surgery, and radiation, which are often not curative and can result in lifelong adverse side effects^14^. We hypothesize that understanding the cascading epigenetic events associated with the functions of the major fusion oncoproteins will illuminate new modalities for precision therapy. The *PAX3::FOXO1* fusion is more prevalent and correlates with a worse patient outcome than *PAX7::FOXO1*^15,16^. Therefore, there is a significant need to understand in vivo PAX3::FOXO1 mechanisms to further inform targeted therapeutics. Leveraging new animal models catalyzes essential observations to enable understanding context-dependent activities in the complexity of in vivo environments.

PAX3::FOXO1 is known to bind to a composite DNA motif of adjacent paired-box and homeobox motif sequences^11^. It establishes networks of myogenic enhancers and co-localizes with chromatin remodelers to regulate oncogenic gene expression signatures^17–19^. We hypothesized that to facilitate chromatin restructuring at the genome scale, PAX3::FOXO1 would have an intrinsic capacity to recognize a diversity of chromatin environments, including open and closed states. To evaluate this hypothesis, our group previously showed that PAX3::FOXO1 has pioneering activity and can interact with closed chromatin, in addition to accessible DNA, in patient-derived cell lines and an inducible human myoblast model^20^. Pioneer factors are a distinct class of transcription factors that can uniquely bind sites occluded by nucleosomes or heterochromatin, in addition to euchromatin binding, to alter the chromatin landscape and inform cell-fate decisions^21,22^. Pioneer factors can function within nucleosomal DNA to promote chromatin decompaction and recruitment of chromatin regulatory machinery^23–28^. Since pioneer factors regulate transcriptional signatures and dependencies, we hypothesized that understanding these mechanisms in vivo would reveal targetable vulnerabilities critical for tumor initiation. In FP-RMS, PAX3::FOXO1 is a core dependency and is transforming in genetic mice and zebrafish models^12,29–31^. We previously developed the zebrafish PAX3::FOXO1-driven tumor model, which we utilized to elucidate disease biology. Tumors were consistent with the human disease and allowed us to identify targets with prognostic significance, such as HES3^12^. PAX3::FOXO1 has been robustly investigated during tumor maintenance. Yet, it remains unclear how this fusion oncoprotein initially interacts with and alters chromatin in vivo, nor are the resulting functional consequences understood.

In this study, we developed an embryonic zebrafish model to study the earliest stages of PAX3::FOXO1 activity with a new, quantitative PerCell ChIP-seq spike-in analytical pipeline to precisely identify PAX3::FOXO1 binding sites and changes to the 2D chromatin landscape. With our approach, we show that the human PAX3::FOXO1 oncoprotein is functionally active in zebrafish and binds to closed chromatin through its homeobox domain. PAX3::FOXO1 reorchestrates the chromatin landscape to promote neural transcriptional signatures that recapitulate a neural cluster observed in FP-RMS. Overall, our system provides insight into the in vivo mechanisms and immediate chromatin regulatory functions of PAX3::FOXO1. Our approach provides a broadly implementable framework to define initial oncogenic and pioneering activities with comparative and quantitative analysis.

## Results

### Establishment of an mRNA injection zebrafish model to study initial PAX3::FOXO1 in vivo activities

We developed a robust high-throughput mRNA injection approach in embryonic zebrafish to investigate the functions of human fusion oncoproteins. We injected single-cell wildtype zebrafish embryos with an mRNA construct containing either the human PAX3::FOXO1 (P3F) coding sequence or the control (CNTL) backbone **(Fig. 1a)**^7^. We cloned the PAX3::FOXO1 coding sequence into the CNTL-2A-sfGFP backbone (adapted from Prykhozhij et al., 2017, ref.^32^) and converted it to mRNA. This construct illuminates fusion oncoprotein expression with GFP fluorescent visualization and a self-cleaving viral 2A to maintain independent protein function^33^. Our model enables a cell agnostic approach to assess PAX3::FOXO1 developmental activity in all three germ layers, which is a powerful approach given the unclear nature of the tumor cell of origin **(Fig. 1b**, ref.^34–36^).

**Fig. 1:**
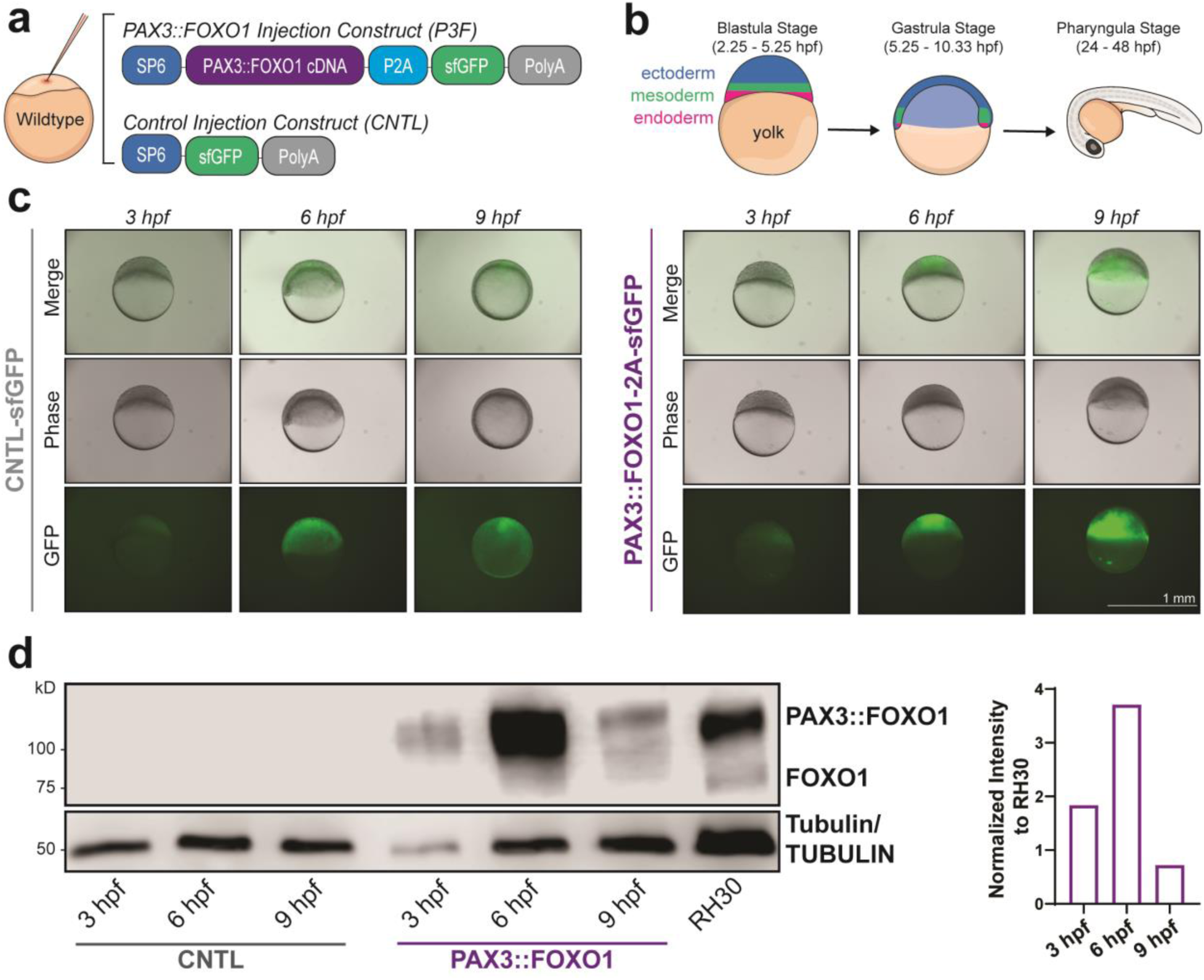
Zebrafish mRNA injection model highly expresses human PAX3::FOXO1 during embryonic development. (a) Wildtype zebrafish embryos were injected at the one-cell stage with 100 ng/μL of PAX3::FOXO1 (P3F) or equal molarity of control (CNTL) mRNA. (b) Schematic of embryonic zebrafish development with positioning of primary germ layers at key time points used in experiments. (c) Representative merged, phase contrast, and GFP images of zebrafish embryos at 3, 6, and 9 developmental hours-post-fertilization (hpf). Scale bar, 1 mm. (d) Representative western blot with protein lysate from 12 embryos per lane for each condition and 8 μg of protein lysate from RH30 cells, a PAX3::FOXO1-positive patient-derived cell line. The PAX3::FOXO1 time-course was repeated for a total of four replicates. Normalized quantification for the representative western blot of PAX3::FOXO1/αTUBULIN signal compared to RH30 cells is on the right, n=1.

To assess the kinetics of PAX3::FOXO1 expression, we evaluated GFP fluorescence (a proxy for PAX3::FOXO1) and PAX3::FOXO1 protein levels. We injected wildtype embryos, imaged and quantified GFP expression, and observed a time-dependent increase in GFP expression from 3 to 9 hours post-fertilization (hpf) **(Fig. 1c**; **Extended Data 1a)**. GFP flow cytometry showed higher fluorescence at 6 hpf versus 12 hpf, suggesting a potential drop-off in translation at later time points **(Extended Data 1b-d)**. We then directly evaluated human PAX3::FOXO1 protein levels by western blot and observed peak expression at 6 hpf before rapidly decreasing by 9 hpf **(Fig. 1d)**. Importantly, we used an antibody specific to the C-terminus of human FOXO1 to detect the PAX3::FOXO1 fusion oncoprotein, which did not cross-react to endogenous zebrafish Foxo1 **(Extended Data 1e-f)**. Given this consistent and high level of PAX3::FOXO1 expression at 6 hpf, we focused on this time point to understand the fusion oncoprotein’s initial activities.

PAX3::FOXO1 expression resulted in phenotypic consequences, which included near complete lethality by 24 hpf and a developmental arrest phenotype prior to gastrulation **(Extended Data Fig. 2)**. Moreover, these phenotypes were consistent with titrating concentrations (25, 50, and 100 ng/µL) of PAX3::FOXO1 mRNA **(Extended Data Fig. 3)**. Since no phenotypic differences were observed across mRNA concentration, we decided to inject 100 ng/µL of PAX3::FOXO1 mRNA to obtain the highest expression of the fusion oncoprotein for all discussed experiments.

### PAX3::FOXO1 interacts with inaccessible chromatin through its homeobox domain

We implemented our mRNA injection model to understand how PAX3::FOXO1 initially engages chromatin to promote early epigenetic programs **(Fig. 2a)**. First, we confirmed PAX3::FOXO1 associated with chromatin through histone extraction experiments. Interestingly, PAX3::FOXO1 was precipitated with histones, as are the pioneer factors Pou5f3 (zebrafish ortholog to OCT4) and Nanog, while the non-pioneer factor Mycn lacked this interaction with the core components of nucleosomes **(Fig. 2b**; **Extended Data Fig. 4a)**. To describe chromatin binding mechanisms of PAX3::FOXO1, we determined its genomic localization through ChIP-seq with the FOXO1-specific antibody (that detects PAX3::FOXO1) from **Figure 1d** (ref.^20^). We applied a PerCell ChIP-seq approach with our zebrafish samples (see companion manuscript, Tallan et al., 2024). Here, we used RH30 cells, a patient-derived PAX3::FOXO1-positive cell line, for spike-in normalization with our zebrafish cells. This data was analyzed through a PerCell ChIP-seq pipeline specifically designed for the analysis of an orthologous cellular spike-in approach, conceptually related to AQuA-HiChIP, for highly quantitative protein-genome binding quantification across experimental conditions^37^. We performed differential binding analysis between anti-FOXO1 ChIPs in our negative CNTL-injected embryos, which lacked the fusion oncoprotein, and PAX3::FOXO1-injected embryos to account for non-specific antibody binding. Our analysis identified 6,564 high-confidence PAX3::FOXO1 binding sites that were consistent between replicates **(Fig. 2c**; **Extended Data 4b)**. Annotation of these high-confidence PAX3::FOXO1 sites showed binding predominantly occurred at distal intergenic or intronic regions, with considerable genomic distance from transcriptional start sites **(Fig. 2d**, **Extended Data 4c)**. This finding suggests that in our zebrafish model, PAX3::FOXO1 primarily binds to regions that are ultimately primed for enhancer function. PAX3::FOXO1’s capacity to bind active enhancers has been noted in the literature in patient-derived tumor cell lines^11,17,20^. However, we sought to understand how initial targeting or DNA binding site selection may occur in a diversity of in vivo epigenomic environments.

**Fig. 2:**
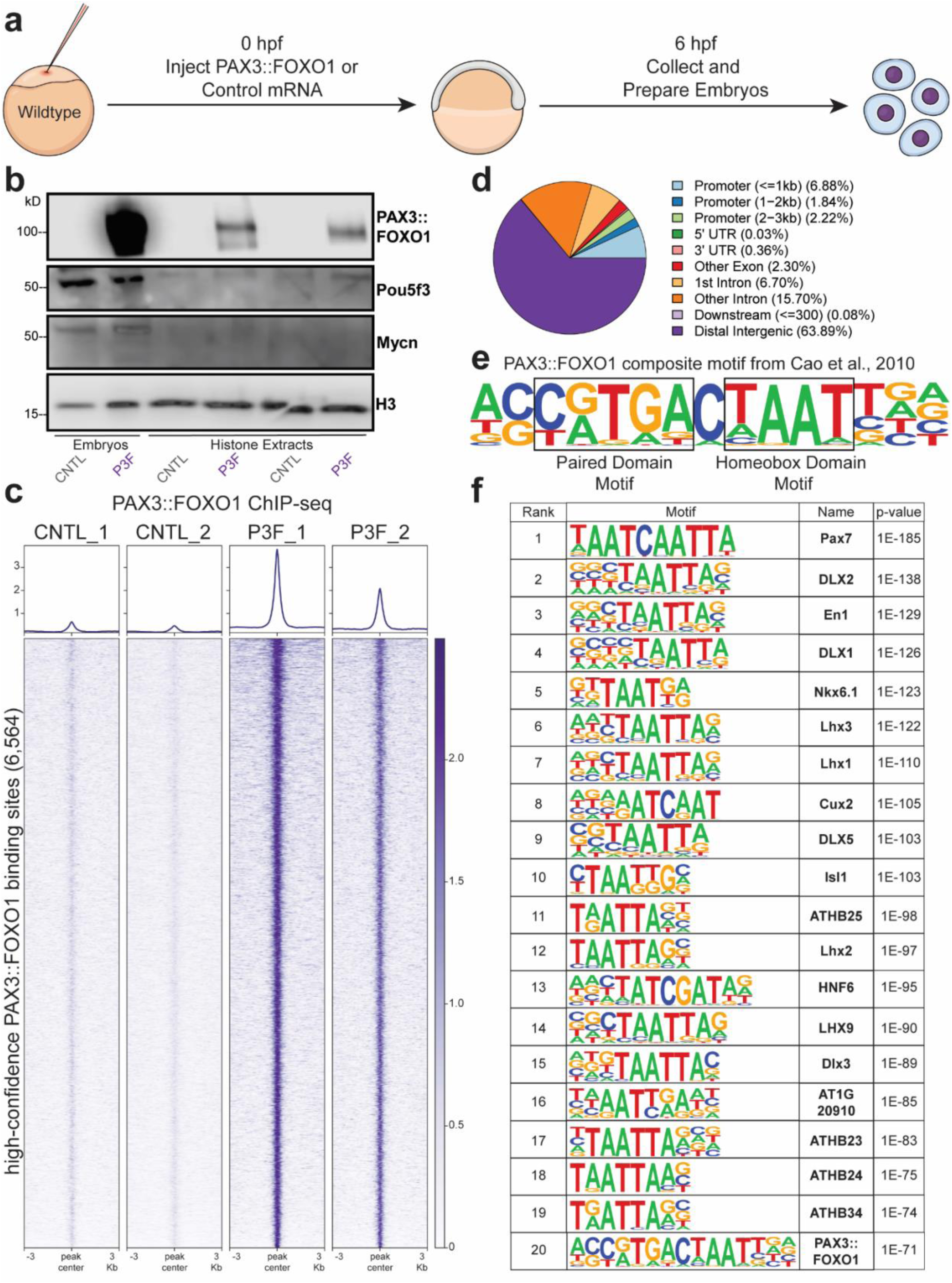
In vivo PAX3::FOXO1 ChIP-seq shows initial binding to homeobox-related motifs. (a) Experimental schematic of mRNA injection with samples collected at the 6 hours post-fertilization (hpf) developmental time point. (b) Western blot on whole embryos (far left two) and histone extractions for protein-DNA interaction by blotting for PAX3::FOXO1, Pou5f3 (pioneer transcription factor), Mycn (non-pioneer transcription factor), and histone H3 in control-injected (CNTL) and PAX3::FOXO1-injected (P3F) embryos. This experiment was repeated for six replicates across three injection days. (c) PAX3::FOXO1 ChIP-seq signal across CNTL and P3F replicates at 6,564 high-confidence PAX3::FOXO1 binding sites. PAX3::FOXO1 binding sites were identified by MACS2 and had significantly different enrichment in P3F versus CNTL samples (edgeR, p=0.05), see methods for additional peak determination information. PAX3::FOXO1 ChIP-seq was completed in duplicate from different injection days. (d) Annotation of PAX3::FOXO1 binding sites based on genomic localization. (e) Schematic of the composite PAX3::FOXO1 motif, which was originally identified by Cao et al., 2010^11^. (f) Top 20 most enriched known motifs at PAX3::FOXO1 binding sites by HOMER.

The known composite PAX3::FOXO1 motif contains its paired and homeobox domain sequences **(Fig. 2e)**. Previously, we showed that initial PAX3::FOXO1 binding sites in human myoblasts are enriched for degenerate bZIP and bHLH motifs^20^. To delineate its mechanisms of in vivo DNA-binding site selection, we performed HOMER motif analysis at our high-confidence PAX3::FOXO1 binding sites and observed enrichment for homeobox domain-containing motifs **(Fig. 2f)**. Of note, our most enriched motif was a PAX7 binding motif that contains a homeobox and inverted homeobox motif (TAAT; ref.^38^). The vast majority of the most enriched motifs (17/20) contained a homeobox domain sequence, while the composite (paired box/homeobox) PAX3::FOXO1 motif was the 20th ranked motif. This motif analysis suggests PAX3::FOXO1 predominantly utilizes partial motif recognition through its homeobox domain to initially interact with DNA instead of binding to its composite motif.

Since PAX3::FOXO1 was associated with histones (as are other pioneer factors), we further investigated its potential pioneering activity in vivo. We examined genomic accessibility by performing ATAC-seq to provide epigenomic context for our model. By overlaying ATAC-seq signal at PAX3::FOXO1 binding sites, we observed that PAX3::FOXO1 robustly enhances DNA accessibility upon its binding **(Fig. 3a)**. Interestingly, when we visualized accessibility with k-means clustering, cluster 3 had minimal accessibility in CNTL-injected embryos with only a modest increase with PAX3::FOXO1 **(Fig. 3b)**. This data suggested PAX3::FOXO1 was bound to lowly accessible chromatin. We found that the majority of PAX3::FOXO1 binding sites were initially inaccessible in control embryos. However, after PAX3::FOXO1 injection, 1,460 of the PAX3::FOXO1 binding sites remained in inaccessible chromatin, which did not overlap with an ATAC-seq peak **(Fig. 3c)**. We validate that these peaks have a strong PAX3::FOXO1 binding signal and no ATAC-seq signal **(Fig. 2d-e**; **Extended Data Fig. 4d-e)**. Two representative tracks are shown for PAX3::FOXO1 binding at *dmd* and *gas7a* **(Fig. 2f**; **Extended Data Fig. 4f)**. PAX3::FOXO1 binding sites in inaccessible chromatin were enriched for homeobox motifs, leading us to hypothesize that PAX3::FOXO1 utilizes its homeobox domain for partial motif recognition at inaccessible chromatin **(Fig. 3c)**. In an inducible human myoblast system, PAX3::FOXO1 has a higher binding intensity at 8 hours post induction when it is sampling DNA motifs as compared to 24 hours post induction when it binds to its composite DNA motif sites^20^. Therefore, we anticipated that stronger PAX3::FOXO1 binding intensity would occur at sites where PAX3::FOXO1 was sampling inaccessible chromatin. We identified PAX3::FOXO1-bound homeobox-rich motifs (i.e., the highly-enriched PAX7 motif) and composite motifs. Homeobox motifs had a higher signal intensity of PAX3::FOXO1 binding than the composite motif, matching the binding kinetics from the PAX3::FOXO1 inducible myoblast system **(Fig. 3g)**. Lastly, since PAX3::FOXO1 expression could be detected at 3 hpf in injected embryos, we assessed the accessibility of PAX3::FOXO1 binding sites from published ATAC-seq datasets across early zebrafish development^39,40^. The majority of PAX3::FOXO1 binding sites remained inaccessible up to 8 hpf, suggesting that binding to these sites during early zebrafish development would require interacting with closed chromatin **(Fig. 3h**; **Extended Data Fig. 4g-h)**. Overall, this supports that PAX3::FOXO1 can invade inaccessible chromatin in a vertebrate model and does so through partial motif recognition of the homeobox motif.

**Fig. 3:**
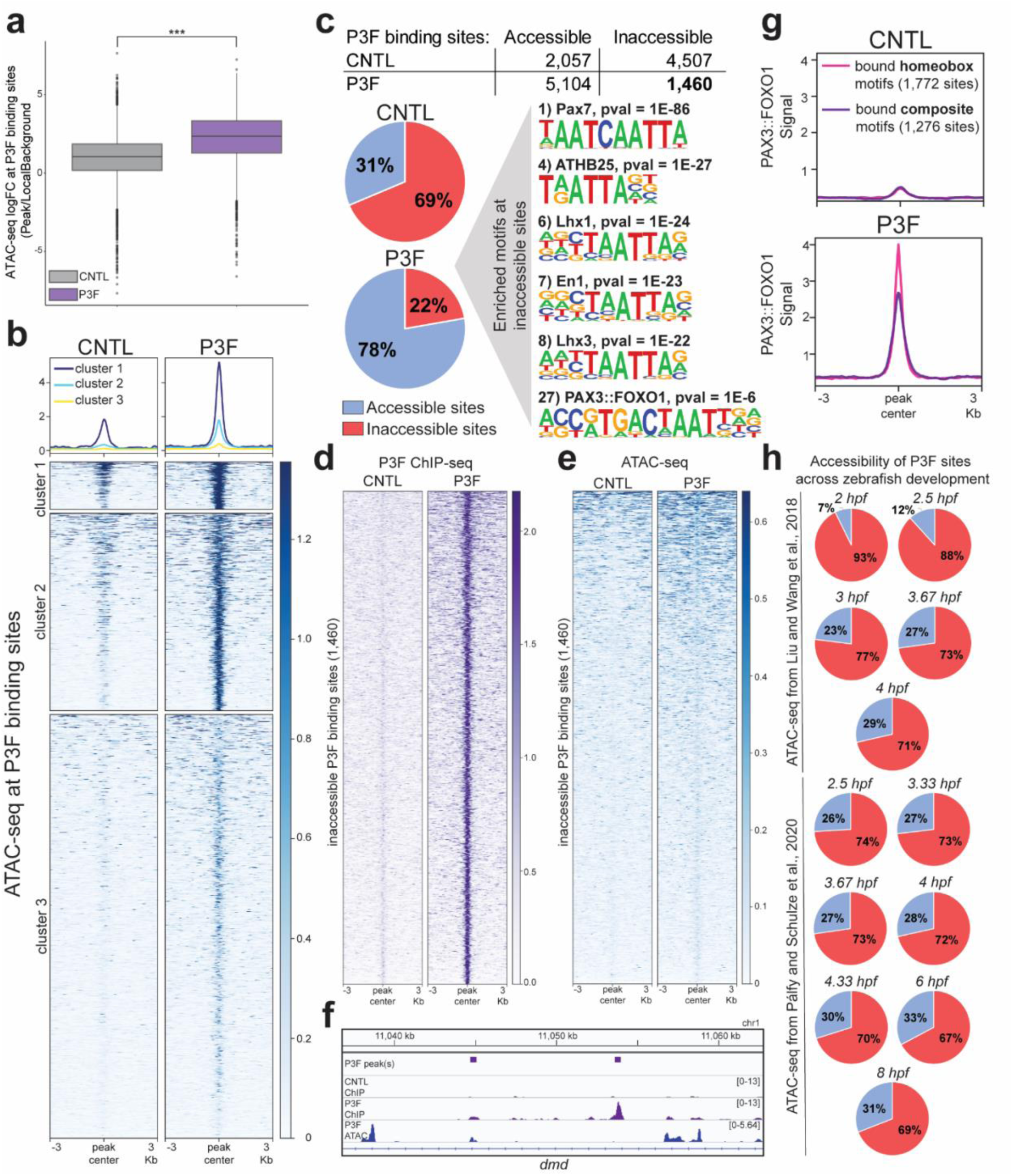
PAX3::FOXO1 invades inaccessible chromatin. (a) Average quantified ATAC-seq signal between replicates at PAX3::FOXO1 binding sites in control-injected (CNTL) and PAX3::FOXO1-injected (P3F) embryos. Line represents the mean. ATAC-seq was completed in duplicate from different injection days. A paired-t-test between conditions was used to determine statistical significance. (b) Visualization of ATAC-seq signal at PAX3::FOXO1 binding sites across conditions. Average ATAC-seq signal between replicates was plotted and stratified by k-means clustering (k=3). (c) Overlap between PAX3::FOXO1 binding sites and ATAC-seq peaks in CNTL and P3F embryos. Accessible sites are defined by binding overlap with an ATAC-seq peak with inaccessible sites lacking an ATAC-seq peak. Known HOMER motif analysis was completed at inaccessible PAX3::FOXO1 binding sites in P3F embryos. (d, e) Average PAX3::FOXO1 ChIP-seq and ATAC-seq signal between replicates at the 1,460 inaccessible PAX3::FOXO1 binding sites, respectively. (f) Representative Integrative Genomics Viewer (IGV) tracks of intronic PAX3::FOXO1 binding in *dmd* that lacks accessibility (right peak). (g) Average PAX3::FOXO1 ChIP-seq signal between replicates at bound homeobox motif binding sites, represented by the Pax7 motif in Fig. 3c, and bound composite motif binding sites, represented by the PAX3::FOXO1 motif in Fig. 3c. Profile plot represents average signal across homeobox and composite motifs, respectively. (h) Overlap between PAX3::FOXO1 binding sites and ATAC-seq peaks throughout early zebrafish development by hours post-fertilization (hpf) from publicly available ATAC-seq data from Liu and Wang et al., 2018^39^ and Pálfy and Schulze et al., 2020^40^. *** denotes p<0.001.

### Homeobox and composite DNA motif preference and binding defines epigenetic roles for PAX3::FOXO1

To delineate the regulatory consequences of PAX3::FOXO1 genomic binding, we compared global chromatin accessibility in our injected embryos to early zebrafish development. While the CNTL-injected embryos clustered near the developmental time course, specifically with gastrulating embryos (~6-8 hpf), our PAX3::FOXO1-injected embryos had a unique chromatin accessibility profile that was distinct from a normal developmental trajectory **(Fig. 4a**; **Extended Data 5a)**. Compared to our CNTL-injected embryos, PAX3::FOXO1 injection resulted in increased accessibility at 23,025 sites and a decrease in accessibility at 17,521 sites **(Fig. 4b)**. Motif analysis at these sites revealed enrichment for the composite PAX3::FOXO1 motif where accessibility increased, suggesting PAX3::FOXO1 was driving these changes, while SOX family members and other developmental transcription factor motifs were enriched where accessibility decreased **(Fig. 4c**; **Extended Data 5b)**. We hypothesized that loss of DNA accessibility at SOX family binding sites might be a signal of PAX3::FOXO1’s ability to systematically alter normal developmental transcriptional programs. Next, we investigated changes in H3K27ac, which marks active chromatin, with our PerCell ChIP-seq approach^41^. H3K27ac deposition was uniquely localized, with most peaks distinct to PAX3::FOXO1-injected embryos **(Fig. 4d**; **Extended Data 5c)**. Differential binding analysis revealed that more sites were activated than repressed (13,863 versus 3,313, respectively), with similar motif enrichments to the ATAC-seq analysis **(Fig. 4e-f**; **Extended Data 5d)**. These analyses suggest that PAX3::FOXO1 generates a distinct chromatin landscape in zebrafish embryos.

**Fig. 4:**
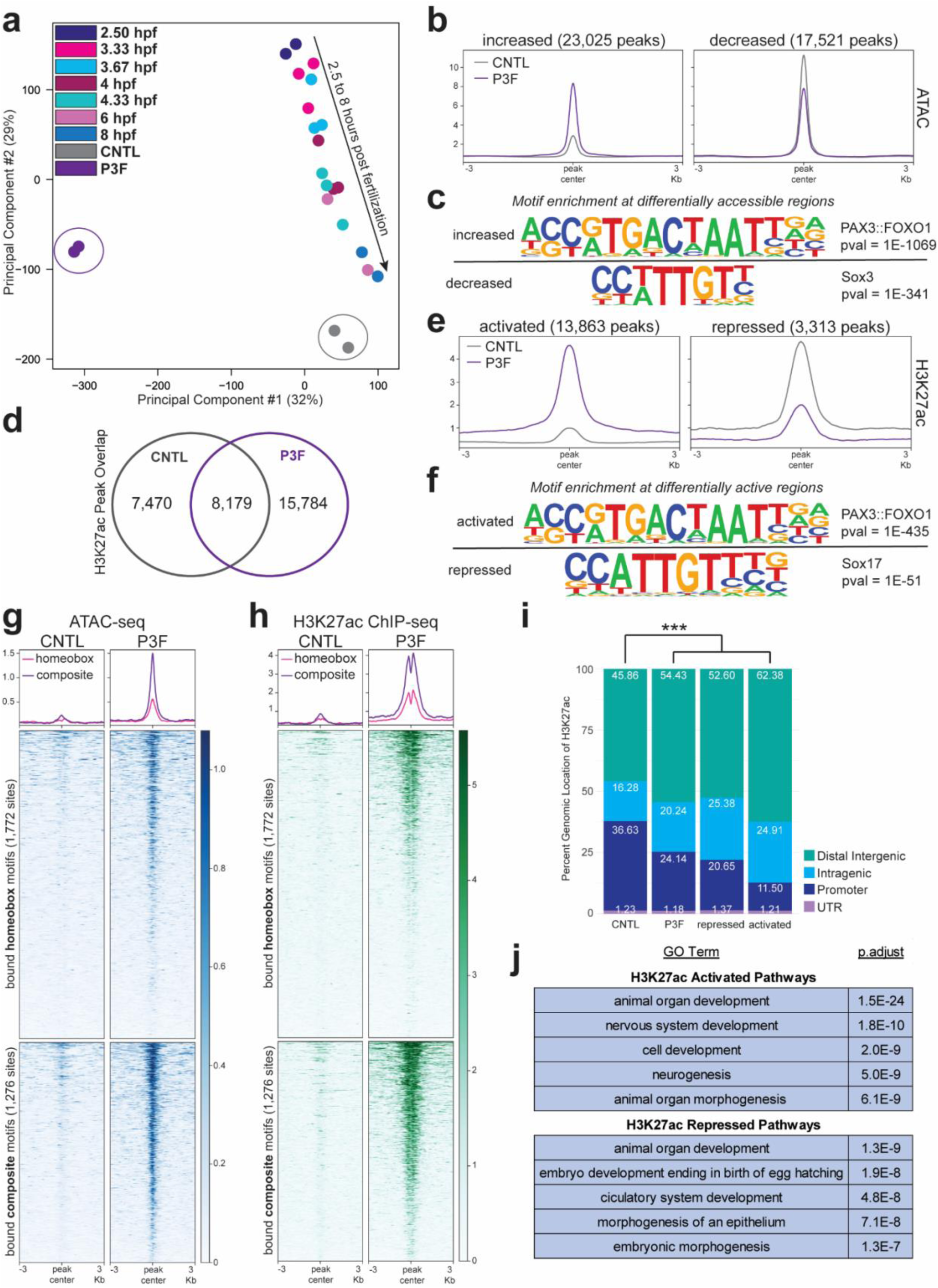
Generation of a neomorphic chromatin landscape is driven by PAX3::FOXO1 binding to its consensus motif. (a) Principal component analysis (PCA) at differentially accessible ATAC-seq peaks in PAX3::FOXO1-injected (P3F) embryos versus control-injected (CNTL) embryos compared to publicly available ATAC-seq from Pálfy and Schulze et al., 2020^40^ across early zebrafish development according to hours post-fertilization (hpf). (b) ATAC-seq signal at sites with increased accessibility (left) and decreased accessibility (right) in P3F versus CNTL embryo. (c) Most enriched known HOMER motif at sites with increased (top) or decreased (bottom) accessibility as shown in Fig. 4b. (d) Overlap of consensus H3K27ac ChIP-seq peaks between replicates in CNTL and P3F embryos. H3K27ac ChIP-seq was completed in duplicate with CNTL embryos from different injection days and P3F embryos from the same injection day. (e) Average H3K27ac ChIP-seq signal between replicates at sites with significantly higher (left) and lower (right) H3K27ac deposition. (f) Most enriched known HOMER motif at sites with higher (top) or lower (bottom) H3K27ac ChIP-seq signal as shown in Fig. 4e. (g, h) Average ATAC-seq and H3K27ac ChIP-seq signal, respectively, at bound PAX3::FOXO1 homeobox or composite motif sites found in Fig. 3g. (i) Annotation of H3K27ac ChIP-seq peaks according to genomic localization in CNTL embryos, P3F embryos, sites with lower H3K27ac in P3F embryos, and sites with higher H3K27ac in P3F embryos, respectively. A Fisher’s exact test with a Bonferroni correction for multiple comparisons on proportion of peaks in promoter regions was used to determine statistical significance. (j) The activity-by-contact model from Fulco and Nasser et al., 2019^42^ inferred which genes were regulated by differentially enriched H3K27ac ChIP-seq peaks. The top 5 enriched gene ontology biological pathways with DAVID analysis for genes regulated by sites with higher H3K27ac in P3F embryos (top) and lower H3K27ac in P3F embryos (bottom). *** denotes p<0.001.

Excitingly, by comparing motif ranks, we observed that, while PAX3::FOXO1’s binding sites were enriched for homeobox sequences, sites with increased accessibility or H3K27ac signal were enriched for the composite PAX3::FOXO1 motif **(Extended Data Fig. 5e)**. We plotted ATAC-seq and H3K27ac signal at PAX3::FOXO1-bound homeobox and composite DNA motifs and observed a greater increase in signal at composite motif sites **(Fig. 4g-h)**. We further investigated how PAX3::FOXO1 modulates H3K27ac and observed that PAX3::FOXO1 injection did not drive a significant increase in overall H3K27ac levels **(Extended Data Fig. 5f-g)**. However, peak annotation revealed that the vast majority of activated sites were at distal intergenic regions **(Fig. 4i)**. Our findings suggest that PAX3::FOXO1 binding to its composite DNA motif is required to drive changes to chromatin and generate active enhancers.

To understand how chromatin binding mechanisms drive gene expression, we identified differentially regulated genes with the activity-by-contact model to determine enhancer-promoter interactions^42^. This model utilizes genomic distance to promoter and enhancer activity from a combination H3K27ac ChIP-seq and ATAC-seq signal to predict enhancer-gene pairs more accurately than using nearest genes. We focused on enhancers with differential levels of H3K27ac from **Figure 4e**, representing regions of chromatin altered by PAX3::FOXO1 activity. Genes regulated by activated enhancers were involved in neural and developmental pathways, while genes regulated by repressed enhancers were enriched for earlier and more general developmental processes **(Fig. 4j)**. Similar pathways were observed with nearest gene analysis **(Extended Data Fig. 5h-i)**. Taken together, these data support that PAX3::FOXO1 binding at its composite DNA motifs promotes chromatin accessibility and histone acetylation, inappropriately activating neural gene targets outside of their normal developmental window.

### PAX3::FOXO1 activates neural and rhabdomyosarcoma transcriptional signatures in zebrafish embryos

To further link chromatin alterations to transcriptional outputs, we completed bulk RNA-seq and compared the transcriptional profiles of PAX3::FOXO1 and CNTL-injected embryos to wildtype zebrafish during blastulation (4.33 hpf) and gastrulation (5.25–8 hpf)^43^. Comparing PAX3::FOXO1-injected embryos to a time course of normal zebrafish development highlighted that the fusion oncoprotein directed a distinct transcriptional signature, while control embryos clustered near gastrulating embryos **(Fig. 5a**; **Extended Data 6a)**. We identified 7,891 (4,270 upregulated, 3,519 downregulated) differentially expressed genes **(Fig. 5b)**. Gene over-representation analysis to published rhabdomyosarcoma gene sets showed that known rhabdomyosarcoma genes were differentially expressed in our model **(Fig. 5c)**. Notable upregulated genes included known PAX3::FOXO1 cooperating genes (*her3*, the zebrafish *HES3* ortholog, *mycn*, and *foxf1*)^12,17,44–46^, a fusion-positive rhabdomyosarcoma-specific histological marker (*olig2*)^47,48^, and genes that inhibit myogenic differentiation in rhabdomyosarcoma (*twist1a* and *six1a*)^49,50^.

**Fig. 5:**
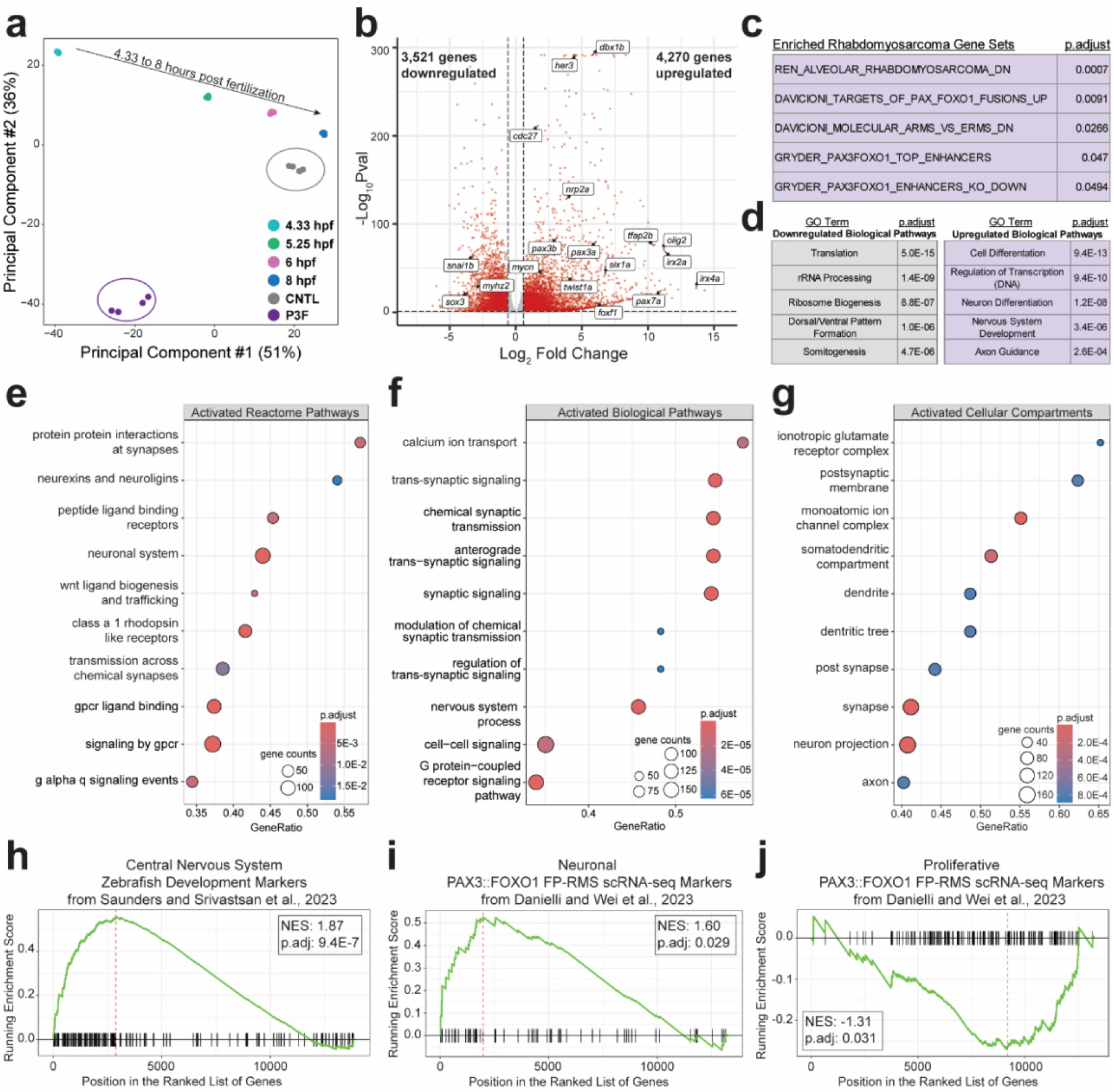
Neural and rhabdomyosarcoma signatures are transcriptionally activated in PAX3::FOXO1-injected embryos. (a) PCA on normalized RNA-seq reads in control-injected (CNTL) versus PAX3::FOXO1-injected (P3F) embryos compared to publicly available RNA-seq during zebrafish blastulation and gastrulation according to hours post-fertilization (hpf) from White et al., 2017^43^. (b) Volcano plot of differentially expressed genes in P3F versus CNTL embryos. Significant genes (red) had an adjusted *p*-value <0.05 and average fold change ≥ 0.586. Highlighted are manually selected genes related to rhabdomyosarcoma, tumorigenesis, myogenesis, and neural development. (c) Enriched rhabdomyosarcoma gene sets from the Molecular Signatures Database (MSigDB) from gene list pathway analysis with differentially expressed genes in P3F embryos. (d) The top 5 enriched gene ontology biological pathways with DAVID analysis for downregulated (left) and upregulated (right) genes in P3F embryos. (e) Gene set enrichment analysis (GSEA) for upregulated pathways in the Reactome pathway database on MSigDB. (f) GSEA for upregulated gene ontology biological pathways. (g) GSEA for upregulated gene ontology cellular compartments. (h) GSEA to markers of the developing zebrafish central nervous system, identified from scRNA-seq analysis from Saunders and Srivatsan et al., 2023^51^. (i-j) GSEA to PAX3::FOXO1-driven rhabdomyosarcoma cell population markers identified from scRNA-seq analysis in Danielli and Wei et al., 2023^52^.

DAVID and gene set enrichment (GSEA) analysis revealed transcriptional repression of ribosomal and developmental processes like somitogenesis, in agreement with the arrested development phenotype seen upon PAX3::FOXO1 injection **(Fig. 5d**; **Extended Data Fig. 6b-c)**. Interestingly, upregulated pathways were strongly enriched for synaptic signaling and receptor-ligand interactions **(Fig. 5d-f)**. Neural-related cellular compartments and molecular functions were also activated, including dendrites, synapses, and various signaling receptors **(Figure 5g**; **Extended Data Fig. 6d)**. This neural activation was distinct to PAX3::FOXO1 injection as these pathways were not prevalent when comparing wildtype late-blastula to gastrulating embryos **(Extended Data Fig. 6e)**. Given this neural pathway activation, we anticipated that PAX3::FOXO1 was driving these cells to a neural state. To characterize the predominant cellular transcriptional identity of these embryos, we utilized embryonic zebrafish tissue markers and observed that the central nervous system was the most enriched tissue type **(Fig. 5h)**^51^. Other enriched tissue types included the peripheral nervous system, eye, and muscle tissues, while liver tissue was repressed **(Extended Data Fig. 6f-h)**. This pattern was consistent when we compared our data to other published cell states as neural cell states were activated and stem cell signatures were repressed **(Extended Data Fig. 6i-j)**. This analysis revealed that the presence of PAX3::FOXO1 initially causes neural transcriptional signature activation in vivo.

Single-cell RNA-sequencing of cell lines, orthotopic patient-derived xenograft models, and patient samples have shown that fusion-positive rhabdomyosarcoma consists of various cellular populations, including differentiated, ground, neuronal, progenitor, and proliferative^52–56^. To assess which transcriptional tumor cell populations PAX3::FOXO1 is initially activating, we completed gene set enrichment analysis to the markers of each of these cell populations. We discovered enrichment of the neuronal cell state and repression of the proliferative state **(Fig. 5i-j)**. This same enrichment pattern is seen in response to chemotherapeutic treatment in mice^52^. Altogether, these findings indicate that in our model, PAX3::FOXO1 initially activates neural signatures, including the neuronal population that is observed in FP-RMS.

### PAX3::FOXO1 regulates gene targets through modulation of the enhancer landscape

We integrated our datasets to understand how PAX3::FOXO1 binding directly impacted chromatin and transcriptional outputs. We focused on PAX3::FOXO1-bound sites that co-localized with H3K27ac deposition. Gene targets of these PAX3::FOXO1-bound enhancers were determined through the activity-by-contact model, as in **Figure 4j**, and were differentially expressed from RNA-seq **(Fig. 6a)**. In this gene set, 1,668 genes were upregulated, 917 genes were downregulated, and the upregulated genes had a more significant change in the magnitude of expression **(Fig. 6b)**. Overall, we discovered 2,585 high-confidence targets that PAX3::FOXO1 initially regulated in vivo.

**Fig. 6:**
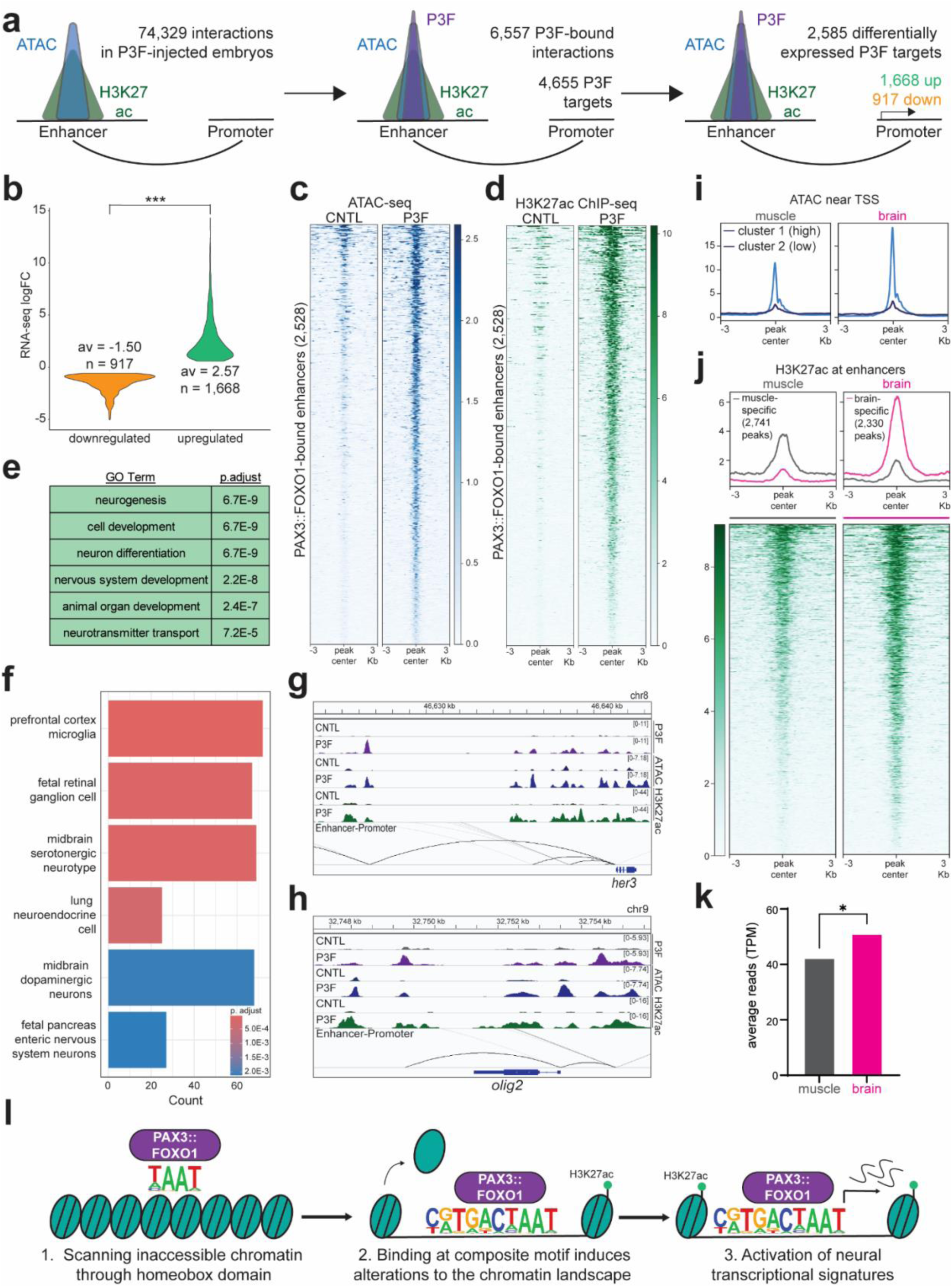
PAX3::FOXO1 directly and epigenetically activates neural-related gene targets. (a) Diagram representing the focus on 2,585 genes that are differentially expressed according to RNA-seq and regulated by a PAX3::FOXO1-bound enhancers in PAX3::FOXO1-injected (P3F) embryos according to the activity-by-contact model. (b) Log fold change expression in P3F versus control-injected (CNTL) embryos of differentially expressed genes in Fig. 6a. A Mann-Whitney test on absolute values was used to determine statistical significance. (c, d) Average ATAC and H3K27ac signal between replicates at the 2,528 PAX3::FOXO1-bound enhancers that regulate activated genes, respectively. (e) The top 5 enriched gene ontology biological pathways with DAVID analysis for genes directly upregulated by PAX3::FOXO1, n=1,668. (f) Top 6 enriched cell type signatures from the Molecular Signatures Database (MSigDB) from gene list pathway analysis with genes directly upregulated by PAX3::FOXO1. (g-h) Representative IGV tracks of activation of neural zebrafish transcription factors *her3* and *olig2* in P3F embryos, respectively. Top tracks are PAX3::FOXO1 ChIP-seq (purple), middle tracks are ATAC-seq (blue), and lower tracks are H3K27ac ChIP-seq (green). Black lines indicate predicted enhancer-promoter interactions for *her3* or *olig2*. (i) ATAC-seq signal at the transcriptional start sites (TSS) of genes directly-upregulated by PAX3::FOXO1 in adult zebrafish muscle and brain tissue from Yang and Luan et al., 2020^62^. Average ATAC-seq signal between replicates was plotted and stratified by k-means clustering (k=2). (j) The activity-by-contact model inferred adult zebrafish muscle and brain-specific enhancers regulating genes directly upregulated by PAX3::FOXO1. Average H3K27ac ChIP-seq seq signal between replicates and across peaks was plotted from Yang and Luan et al., 2020^62^. Heatmaps show average H3K27ac signal between replicates at muscle-specific enhancers in muscle samples (gray) and at brain-specific enhancers in brain samples (pink). (k) Average RNA-seq expression, by TPM, of genes directly upregulated by PAX3::FOXO1 in adult zebrafish muscle and brain tissue from Yang and Luan et al., 2020^62^. A Wilcoxon matched-pairs signed rank test was used to determine statistical significance. (l) Proposed model of initial in vivo activities of PAX3::FOXO1. *** denotes p<0.001 and * denotes 0.01<p<0.05.

In agreement with our previous observation that the majority of H3K27ac sites were unique upon PAX3::FOXO1 injection in **Figure 4d**, PAX3::FOXO1-bound enhancers that directly activated gene targets were specifically active in PAX3::FOXO1-injected embryos **(Fig. 6c-d)**. Given these findings and the established role of PAX3::FOXO1 as a transcriptional activator, we were motivated to investigate how PAX3::FOXO1 directly repressed gene targets. These repressed targets were involved in ribosomal and early developmental processes **(Extended Data Fig. 7a)**. Even though the enhancers that initially regulated these genes lost activity, as demarcated by a decrease in H3K27ac, the PAX3::FOXO1-bound enhancers still gained activity **(Extended Data Fig. 7b-c)**. Globally, we observed that PAX3::FOXO1 binding rarely co-localized with sites that lost H3K27ac, and the majority of its binding sites gained deposition of this histone mark, suggesting PAX3::FOXO1 was not driving heterochromatin formation at these sites **(Extended Data Fig. 7d)**. Since enhancer activity is inversely related to genomic distance, we hypothesized that PAX3::FOXO1 may shift H3K27ac to enhancers farther from repressed genes to drive their repression^57,58^. In support of this hypothesis, we saw that PAX3::FOXO1-bound enhancers were farther away (in kb) from the repressed genes versus the activated ones **(Extended Data Fig. 7e)**. Counterintuitively, these repressed genes had more enhancers regulating them upon PAX3::FOXO1 injection **(Extended Data Fig. 7f)**. However, enhancers with increased activity were >50 kb away from repressed genes with a corresponding loss of enhancer number and activity within 10 kb **(Extended Data Fig. 7g-i)**. Together, our analysis suggests that PAX3::FOXO1 generates de novo enhancers, and PAX3::FOXO1 enhancer activation causes a redistribution and loss of H3K27ac at select genes.

### PAX3::FOXO1 directly activates neural gene targets

PAX3::FOXO1 directly activated genes were enriched for neural processes, including neurogenesis and neurotransmitter transport, and had signatures of various neural cells like neurons and ganglions **(Fig. 6e-f)**. We went to identify 130 transcription factors that were direct PAX3::FOXO1 targets with the rationale that: 1) PAX3::FOXO1 contributes to the core-regulatory circuitry of FP-RMS through cooperating with other transcription factors, and 2) DNA transcription was highly enriched in our RNA-seq data^17,59,60^. These 130 transcription factors included endogenous PAX3::FOXO1 fusion partners (*pax3a*, *pax3b*, and *foxo1b*) **(Extended Data Fig. 8)**. Although our model does not incorporate the genetic translocation into the zebrafish genome, binding near fusion-partners further supports that PAX3::FOXO1 activates its own expression^59^. Broadly, these activated transcription factors were enriched in neural processes and often localized in the zebrafish brain **(Fig. g-h; Extended Data Fig. 9)**. This finding suggests PAX3::FOXO1 activates and may cooperate with this set of transcription factors as an initial activity to promote altered core regulatory circuitry in vivo through its early-immediate chromatin targeting functions^60,61^.

Lastly, we compared the 1,668 activated initial PAX3::FOXO1 targets between zebrafish muscle and brain with the hypothesis that these neural targets should be more activate in the brain^62^. We observed that their promoters were more accessible in brain tissue **(Fig. 6i**; **Extended Data 10a)**. Furthermore, through the activity-by-contact model, we identified enhancers that regulated these genes and determined that while shared enhancers had similar activity, brain-specific enhancers were more active than muscle-specific enhancers **(Fig. 6j**; **Extended Data 10b-e)**. Finally, these genes were more highly expressed in the brain versus the muscle **(Fig. 6k)**. Together, our findings reveal that PAX3::FOXO1’s initially activates neural-related gene targets.

## Discussion

In the past 30 years, significant research efforts have determined that PAX3::FOXO1 is a critical driver and dependency of FP-RMS. However, the primary epigenetic targets, mechanisms, and requirements for tumor initiation, maintenance, or metastasis have remained elusive. It follows that the standard of care therapeutic regime, to which FP-RMS patients are often less responsive, has remained largely unchanged. To address these fundamental issues, we investigated the initial activities of PAX3::FOXO1 in a new in vivo context to understand how this chimeric transcription factor interacts with and modulates chromatin. We find evidence supporting that PAX3::FOXO1 has in vivo pioneering activity through utilizing partial motif recognition of homeobox motifs. Alternatively, PAX3::FOXO1 binding to its composite DNA motif drives changes to the chromatin landscape to support neural transcriptional signatures **(Fig. 6l)**.

In cell culture, PAX3::FOXO1 can localize to heterochromatin fractions, asymmetrically flank the repressive H3K9me3 histone mark, and bind within generally repressive B-chromatin compartments, in addition to binding at active enhancers and within more transcriptionally active A-compartments^17,20,63^. We observed a similar pattern in our zebrafish model, with PAX3::FOXO1 invading regions originally and persistently devoid of chromatin accessibility **(Fig. 3)**. After binding, we see a subsequent increase in chromatin accessibility and H3K27ac levels **(Fig. 4)**. Together, these findings support that PAX3::FOXO1 has in vivo pioneer function, binding to closed chromatin to induce changes to the epigenetic landscape in a diversity of contexts. However, across the zebrafish and human myoblast cell models, the mode of initial chromatin recognition and pioneer activity seems to vary. In inducible myoblasts, we find evidence that PAX3::FOXO1 utilizes degenerate bZIP and bHLH motifs for DNA recognition, given the enrichment of those motifs at PAX3::FOXO1 binding sites^20^. In zebrafish, we observe partial motif recognition and enrichment of homeobox motifs **(Fig. 3c)**. It is possible that, depending on context, PAX3::FOXO1 could utilize different binding modes to facilitate interactions with nucleosomal or repressive chromatin. Additional factors may include PAX3::FOXO1 expression levels or timing and the duration of its expression. Even though pioneer factor activity can occur at nucleosomal or inaccessible chromatin, context can influence pioneer factor function as well. The deposition of various histone marks can influence pioneer factor binding, and the availability of cooperating transcription factors can modulate which sites are activated^64–66^. As such, we found that only 1,276 out of 274,499 composite PAX3::FOXO1 motifs in the zebrafish genome are bound by PAX3::FOXO1, suggesting that features in the chromatin landscape of these developing embryos might provide a secondary layer of logic for DNA binding site selection. While identifying direct targets of PAX3::FOXO1, neither *myod1* nor *myog1* were directly activated even though these factors are highly expressed and co-localize with PAX3::FOXO1 at myogenic enhancers in FP-RMS^17,59^. Instead, we observed activation of neural transcription factors and gene targets **(Fig. 6**; **Extended Data Fig. 9)**. Therefore, it could be therapeutically relevant to understand the motif selection process of PAX3::FOXO1 and critical targetable cooperating factors that prevent PAX3::FOXO1 from binding to oncogenic enhancers or influence its activation of oncogenic targets.

We find evidence that PAX3::FOXO1 utilizes the homeobox domain for heterochromatin interactions **(Fig. 3)**. Previous literature supports that the homeobox domain is critical for PAX3::FOXO1-driven tumorigenesis. In a 3T3 mouse embryonic fibroblast system, ectopic expression of PAX3::FOXO1 with mutations in the homeobox domain abrogated anchorage independent growth, whereas point mutations in the paired domain did not^67^. However, a recent CRISPR/Cas9 domain screen in RH4 cells, a PAX3::FOXO1-positive patient-derived cell line, showed that independently mutating both DNA binding domains suppresses 2D cell proliferation, highlighting the importance of the context and assay for interpretation of results^68^. Normally, PAX3 functions as a neural crest transcription factor and utilizes its homeobox domain for mitotic chromosomal loading^69^. Altogether, the homeobox domain is especially critical for PAX3::FOXO1 pioneering and oncogenic activity. These data support the necessity of PAX3::FOXO1 pioneering activity for tumorigenesis.

In our model, we observe that ~22% of PAX3::FOXO1 binding sites remain inaccessible and that ~42% failed to overlap with H3K27ac **(Fig. 3c**; **Extended Data Fig. 7d)**. This shows that, like other pioneer factors, PAX3::FOXO1 binding can precede, and does not always correlate to, gene activation^23,70,71^. Moreover, PAX3::FOXO1 binding to its composite versus the homeobox motif results in more prominent chromatin accessibility and H3K27ac deposition **(Fig. 4g-h)**. This finding suggests a potential biological window where PAX3::FOXO1 may lack regulatory and functional consequences. Drug discovery-based studies may elucidate targetable interactions between PAX3::FOXO1 and chromatin regulators like BRG1, CHD4, and CBP/p300, which contribute to its activities^18,19,68,72^. The pharmacological inhibition of these epigenetic factors could be broadly applicable to fusion-driven sarcomas, given their reliance on chromatin regulatory mechanisms^73,74^. Furthermore, covalent small molecular inhibitors of FOXA1 can modulate FOXA1 pioneer function and could potentially be applied to FP-RMS^75^. Potentially, stalling or re-directing PAX3::FOXO1 binding at homeobox motifs and preventing chromatin remodeling could be a valuable approach to suppress tumorigenesis.

Our model showcases that an initial function of PAX3::FOXO1 is to directly activate neural transcriptional programs **(Fig. 5**; **Fig. 6)**. Even though rhabdomyosarcoma is classically described by myogenic characteristics, we see that FP-RMS patients have neural pathways enriched versus the less-aggressive fusion-negative subtype^48^. Consistent with the neural signatures found in our model, PAX3::FOXO1 electroporation into the neural tube of chicken embryos could transdifferentiate neural cells into cells with rhabdomyosarcoma characteristics^76^. Neuronal populations were also distinctly identified in FP-RMS patient and orthotopic patient-derived xenograft tumor samples from single-cell RNA-sequencing^53^. Critically, upon chemotherapeutic treatment in these xenograft models, this neuronal population was transcriptionally enriched, which would support a hypothesis that neural pathways drive FP-RMS therapeutic resistance^52^. Through our PAX3::FOXO1 zebrafish tumor model, we found that the neural transcription factor *her3* (zebrafish ortholog to *HES3*) was one of the most upregulated genes^12^. Here, we report initial activation of the neuronal and repression of the proliferative cell populations in our model, in concordance with what was seen following xenograft drug treatment **(Fig. 5h-i)**. These similarities suggest two key conceptual integrations of our observations: (**i**) neural signatures may play a role across various stages of FP-RMS tumorigenesis and (**ii**) that therapeutic resistance may be a recapitulation of tumor initiation.

The mRNA injection model we developed allowed us to evaluate the earliest in vivo activities of the PAX3::FOXO1 fusion oncoprotein. This model generates samples in a high-throughput manner and is adaptable to study other fusion oncoproteins and the roles of diverse pioneer factors in a developmental, in vivo context. Comparative analysis of the initial epigenetic mechanisms of various fusion oncoprotein transcription factors like PAX7::FOXO1 in FP-RMS or fusion oncoproteins across pediatric sarcomas could help identify shared or divergent targetable mechanisms. Given that PAX7::FOXO1 is less aggressive than PAX3::FOXO1 clinically, our model could mechanistically define how fusion oncoprotein pioneering/biological activity manifests into divergent clinical outcomes^4,15,16,77^. Further, our model highlights the importance of studying the epigenetic mechanisms of pediatric oncoproteins, or any oncoprotein, in a dynamic developmental context. More broadly, we describe a comparative platform to understand how epigenetic activities can influence the tumorigenic process to identify new oncogenic dependencies.

## Online Methods

The research in this study follows ethical regulations. Research procedures were approved by the IACUC (protocol AR19-00172) at The Abigail Wexner Research Institute at Nationwide Children’s Hospital.

### Zebrafish Husbandry

Zebrafish (*Danio rerio)* were maintained in an AAALAC-accredited, USDA-registered, OLAW-assured facility at Nationwide Children’s Hospital. The facility is in compliance with the Guide for the Care and Use of Laboratory Animals. Adult zebrafish are housed in 0.8-L, 1.8-L, 2.8-L, or 6-L tanks in mix-sex groups with a density of 5-12 fish per liter on a recirculating system (Aquaneering, San Diego, CA) in 28°C water (conductivity, 510 to 600 μS; pH, 7.3 to 7.7; hardness, 80 ppm; alkalinity, 80 ppm; dissolved oxygen, greater than 6 mg/L; ammonia, 0 ppm; nitrate, 0 to 0.5 ppm; and nitrite, 0 ppm). The fish room has a 14:10-h light:dark cycle. Zebrafish are free of *Pseudoloma neurophilia*, *Pleistophora hyphessobryconis*, *Pseudocapillaria tomentosa*, *Mycobacterium spp.*, *Edwardsiella ictalurid*, *Ichthyophthirius multifilis*, *Flavobacterium columnare*, and *zebrafish picornavirus (ZfPV1)*, which was determined by quarterly sentinel monitoring. System water was carbon-filtered (20-μm pleated particulate filter, 40W UV light exposure) from municipal tap water. From 5-30 days post-fertilization, the fish were fed live rotifer feeds three times a day and after 30 days post-fertilization were fed with a commercial pelleted diet twice daily. Zebrafish embryo micro-injections were sex-blinded as zebrafish sex is not determined until 30–60 days of age in our facility. WIK zebrafish were initially obtained from the Zebrafish International Resource Center (ZIRC; https://zebrafish.org/) and used as the wildtype strain in this study.

### Plasmid Cloning and mRNA Injection Construct Generation

The human PAX3::FOXO1 coding sequencing was originally synthesized and cloned from Shapiro et al., 1993^7^. Primers for plasmid cloning and generation are listed in Supplemental Table 1. Restriction digest sites were added to the PCR product. The PCR product was digested with *Pac1* and *Asc1* and cloned into pCS2+MCS-P2A-sfGFP in a 3:1 insert to backbone ratio with T4 DNA ligase (NEB, M0202S). The initial plasmid was a kind gift from Jason Berman (Addgene plasmid #74668, ref.^32^). Ligated plasmids were transformed in DH5α cells (Fisher Scientific, FEREC0111) and validated with Sanger sequencing. The PAX3::FOXO1-2A-sfGFP and pCS2+MCS-P2A-sfGFP plasmids were linearized with *Not1*, ethanol precipitated, and mRNA transcribed from linearized plasmids with the mMESSAGE mMACHINE™ SP6 Transcription Kit (ThermoFisher Scientific, AM1340). An additional CNTL-sfGFP negative control was created by amplifying out the sfGFP and polyA tail of pCS2+MCS-P2A-sfGFP. PCR products underwent A-tailing and ligation with the pGEM-T Easy kit (Promega, A1360). Constructs were transformed, validated, and transcribed to mRNA as described above. Plasmids are in the process of being deposited to Addgene.

### Zebrafish Embryo Injection

Adult wildtype WIK zebrafish were in-crossed. Offspring were injected in the yolk at the single-cell stage with 100 ng/μL of PAX3::FOXO1-2A-sfGFP or equal molarity of the control mRNA (unless otherwise noted) and a drop size diameter of 0.15 mm. Injection mixes consisted of mRNA and 0.05% phenol red in 3X Danieau’s buffer (52.2 mM NaCl, 0.63 mM KCl, 0.36 mM MgSO_4_·7H_2_O, 0.54 mM Ca(NO_3_)_2_·7H_2_O, 4.5 mM HEPES). The embryos were incubated in 1X E3 buffer (5 mM NaCl, 0.17 mM KCl, 0.33 mM CaCl_2_, 0.33 mM MgSO_4_) at 32°C until they reached the desired developmental time point/stage for that temperature according to Kimmel et al., 1995^34^.

### Phenotypic Analysis

Embryonic survival was determined by counting the initial number of injected embryos and the number of dead embryos at each labeled developmental time point. The developmental stage was visually assessed according to known embryo phenotypes as described by Kimmel et al., 1995^34^. GFP fluorescence was measured from embryo images with a Leica M205FA fluorescent stereoscope. Image analysis by raw integrated density quantification of whole embryos was performed in ImageJ.

### Flow Cytometry

Zebrafish embryos were injected at the one-cell stage with injection mixes containing either 50 ng/μL of PAX3::FOXO1 mRNA or 50 ng/μL of pCS2+her3^WT^-P2A-sfGFP mRNA and 50 ng/μL of pCS2-tagRFPT.zf1. pCS2+her3^WT^-P2A-sfGFP and pCS2-tagRFPT.zf1 mRNA were prepared as described in Kent et al., 2023^44^. PAX3::FOXO1 injected embryos were incubated at 32°C until they reached the desired developmental stage of 6 hpf or 12 hpf. pCS2+her3^WT^-P2A-sfGFP injected and uninjected WIK embryos were incubated at 28.5°C and collected at 24 hpf to set the fluorescent gate. For processing, embryos were dechorionated with pronase, deyolked (55 mM NaCl, 1.8 mM KCl, 1.25 mM NaHCO_3_), and washed with 0.5X Danieau’s buffer (29 mM NaCl, 0.35 mM KCl, 0.2 mM MgSO_4_·4H_2_O, 0.3 mM Ca(NO_3_)_2_·4H_2_O, 2.5 mM HEPES). Embryos were dissociated into single cells by passing through a 40 μM cell strainer. Cells were suspended in PBS and fixed in 1% formaldehyde for 10 minutes at room temperature. Fixation was quenched in 0.125M glycine for 5 minutes on ice and washed in PBS before being stored at 4°C until flow cytometry. Flow cytometry was completed by the Nationwide Children’s Hospital Flow Cytometry Core.

### Western Blotting

Embryos were collected at the desired developmental time point. Pools of embryos were dechorionated and deyolked as described above. Embryos were snap-frozen with dry ice and stored at −80°C. Protein lysates were generated by adding 2 μL of 2X Laemmli Buffer (BIO-RAD, 1610737) per embryo and heating samples for 5 minutes at 95°C. Lysates were briefly spun down. Equal volume and embryo number were loaded in a 10% Mini-PROTEAN® TGX Stain-Free™ Protein Gel (BIO-RAD, 4568034). Samples were transferred to a 0.2 μm PVDF membrane (BIO-RAD, 1620177). The membrane was blocked with Casein and 0.05% Tween-20 (Fisher Scientific, PI37528). Following blocking, the membrane was incubated with the appropriate following antibodies at 4°C overnight at 1:1000 concentration in the blocking solution: αFOXO1 (Cell Signaling, C29H4), αTUBULIN (Cell Signaling, 3873S), αPou5f3 (GeneTex, GTX132245), αMycn (ThermoFisher Scientific, PA5-117717), αNanog (GeneTex, GTX132491), αH3 (Abcam ab1791), or H3K27ac (Active Motif, 39133). Membranes were washed with 1X PBS with 0.05% Tween-20 before incubating for 1 hour at room temperature with 1:10,000 of the appropriate secondary antibody in blocking solution: HRP-α-mouse (BIO-RAD, 1706516) or HRP-α-rabbit (BIO-RAD, 1721019). Following another PBS with 0.05% Tween-20 wash, the membranes were imaged on a C-DiGit Chemiluminescent LI-COR imager (VWR, 103375-240) with either SuperSignal West Pico PLUS Chemiluminescent Substrate (Fisher Scientific, PI34577) or SuperSignal West Atto Ultimate Sensitivity Chemiluminescent Substrate (Fisher Scientific, PIA38554).

### Histone Extraction

Embryos were injected with either PAX3::FOXO1-2A-sfGFP or CNTL-sfGFP mRNA, and aliquots of at least 55 embryos were collected, as described above. Histone extraction was adapted from Shechter et al., 2007^78^. Aliquots were thawed and suspended in 800 µL of Buffer A (0.02 M HEPES pH = 7.9, 0.01 M KCl, 0.0002 M EDTA) with proteinase inhibitors. Embryos were dissociated with mechanical mortar and pestle and incubated on ice for 10 minutes. 40 µL of 10% NP-40 in buffer A was added, and the samples were vortexed 2 x 15 seconds with 1-minute ice incubation in between. Samples were centrifuged at 4°C, 10,000g for 1 minute. The supernatant was discarded and the nuclear pellet was re-suspended in 400 µL of 0.4N H_2_SO_4_ in Buffer A. Samples were incubated at 4°C with overhead rotation for at least 2 hours, then centrifuged at 4 °C, 10,000g for 10 minutes. 400 µL of the supernatant was transferred to a clean 1.5 mL tube, and histones were precipitated by adding 100% trichloroacetic acid dropwise to a 25% final concentration. The solution was inverted and incubated on ice for at least 30 minutes. The supernatant was gently aspirated following centrifugation at 4 °C, 10,000g for 10 minutes. Two cold acetone washes were completed with a 4°C, 10,000g 5-minute centrifugation between washes. Histones were air-dried at room temperature for 20 minutes, then re-suspended in 30 µL of H_2_O. Histone extraction was characterized by western blotting as described above, with boiling and loading the entire volume into a 4–15% gradient mini-PROTEAN TGX gel (BIO-RAD, 4561084).

### Chromatin Immunoprecipitation (ChIP-seq)

#### PAX3::FOXO1 Chromatin Immunoprecipitation

ChIP-seq protocol was adapted from Sunkel et al., 2021^20^. Embryos were injected with either PAX3::FOXO1-2A-sfGFP or CNTL-sfGFP mRNA and incubated until they reached the 6 hpf developmental stage. Embryos were dechorionated as above and dissociated by thorough pipetting in dPBS. Cells were fixed in a final concentration of 1% formaldehyde for 10 minutes at room temperature and quenched by adding glycine to a final concentration of 125 mM with a 5-minute incubation on ice. Cells were aliquotted, snap-frozen, and stored at −80°C. For spike-in normalization, PAX3::FOXO1-positive RH30 patient-derived cells RH30 (ATCC, CRL-2061) were fixed and stored like the zebrafish samples following their expansion and lifting with TrypLE Express (ThermoFisher Scientific, 12604013). RH30 cells were authenticated by STR and tested for mycoplasma annually through Genetica Inc a subdivision of LabCorp. ChIP-seq experiments were completed in duplicate with 2 million injected zebrafish cells and 250,000 RH30 cells. Duplicates came from different injection clutches. Cells were thawed, combined, and suspended in 300 µL of TE buffer with protease inhibitors. Samples underwent sonication for 5 minutes (30 seconds on, 30 seconds off) with a Bioruptor Pico (Diagenode, B01060010). 5 µL of sonicated DNA was taken as input DNA and reverse cross-linked with 90 µL H2O, 8 µL 18.5 mg/mL Proteinase K, 4 µL 5M NaCl for 1 hour at 65°C at 400 rpm in a ThermoMixer. DNA was purified with the Qiagen MinElute PCR Purification kit (QIAGEN, 28004) and eluted in 36 µL warm EB buffer. Sonication was confirmed by running 1 μL of input DNA on an E-gel (Invitrogen, G8300). The remaining DNA (280 μL) was adjusted to a final buffer concentration of 1% Triton X-100, 0.1% SDS, 0.1% sodium deoxycholate, and 200 mM NaCl and spun at 4°C, 13,000 rpm for 10 minutes. Supernatent was rotated for 2 hours in a cold room with 4 μL anti-FOXO1 (Cell Signaling, C29H4). Then, 40 µL of Dynabeads Protein G (Fisher Scientific, 10-003-D) were added for continued overnight incubation. Samples were washed on a magnetic rack twice in TE with 0.1% SDS, 0.1% sodium deoxycholate, and 1% Triton X-100, twice in TE with 1% Triton X-100, 0.1% SDS, 0.1% sodium deoxycholate, and 200 mM NaCl, twice in TE with 250 mM LiCl, 0.5% NP-40, 0.5% sodium deoxycholate, and once with TE. DNA reverse-crosslinking was completed with a 65°C overnight incubation in 100 μL TE, 2.5 μL 10% SDS, and 5 μL 18.5 mg/mL Proteinase K. DNA was purified in the same manner as the input DNA immunoprecipitation was confirmed by qPCR with 1 µL of ChIP and input DNA using the primers listed in Supplemental Table 1.

#### H3K27ac Chromatin Immunoprecipitation

Zebrafish cells were collected as described for the PAX3::FOXO1 ChIP-seq but with pCS2+MCS-P2A-sfGFP mRNA for the control samples. Here, 1 million injected zebrafish cells and 1 million RH30 cells were thawed and combined in 800 µL of TE buffer with protease inhibitors. Experiments were performed in duplicate, PAX3::FOXO1-injected embryos came from the same injection clutch, and control-injected embryos from different injection clutches. Cells were sonicated for 25 minutes (30 seconds on, 30 seconds off) with a 30% amplitude on an EpiShear probe sonicator (Active Motif, 53051). 5 µL of DNA was reverse cross-linked overnight at 65°C in 20 μL TE, 1 μL 10% SDS, and 1 μL 18.5 mg/mL Proteinase K to serve as the input DNA. The remaining steps were completed in the same manner as the PAX3::FOXO1 ChIP but with 770 µL used for immunoprecipitation and primary antibody incubation with 4 μL anti-H3K27ac (Active Motif, 39133).

#### ChIP-seq Library Preparation

Sequencing libraries were prepared with 12-14 μL of ChIP DNA or 4 μL of input DNA. End-repair was completed with either the End-It DNA End-Repair Kit (Lucigen, ER81050) for H327ac ChIP-seq or the NEBNext End Repair Module (NEB, E6050S) for PAX3::FOXO1 ChIP-seq. The a-tailing reaction went for 30 minutes at 37°C, 300 rpm in a ThermoMixer with 32 µL of purified DNA from the QIAquick PCR Purification kit, 5 μL 10X Klenow Buffer/NEBuffer 2, 10 μL 1 mM dATP, and 3 μL 5U/μL Klenow Exo. Adapter ligation was completed with 22 μL of purified DNA in EB, 3 μL T4 DNA Ligase Buffer, 2 μL Index PE Adapter Mix, and 3 μL T4 DNA Ligase. PE Adapter Mix was generated in advance by annealing top and bottom multiplex adaptor primers in Supplemental Table 1 to a final concentration of 15 μM with a 95°C, 5 minutes incubation, and slow-cool to room temperature for 120 minutes. Ligation was incubated for 30-60 minutes at room temperature. Ligated products were purified with 1.0X AMPure beads selection (Beckham-Coulter, A63880) and eluted in 23 μL EB. DNA was amplified via qRT-PCR until the reaction plateaued (11.4 μL DNA, 12.5 μL Phusion Tag 2X Master Mix, 0.5 μL unique Illumina i5 primer, 0.5 μL unique Illumina i7 primer, 0.15 μL 100X SYBR Green). PCR cycle was at 98°C for 30 seconds, then repeating at 98°C for 10 seconds, 65°C for 30 seconds, and 72°C for 30 seconds. PCR products were purified with a 1.8X AMPure bead selection and final elution in 23 μL EB. Sequencing was completed by Nationwide Children’s Hospital Institute for Genomic Medicine (IGM) on a NovaSeq SP with 2×150 bp following additional QC by Agilent Bioanalyzer and Qubit.

#### Data analysis

ChIP-seq analysis was automated through the PerCell ChIP-seq pipeline in Tallan et al., 2024. In brief, samples were aligned to danRer11 with Bowtie2^79^. PCR duplicates were removed with PICARD (https://broadinstitute.github.io/picard/). Peaks were called and bigwigs were generated with MACS2^80^. Consensus PAX3::FOXO1 peaks were identified with MACS2 with a *p*-value (Poisson test, ppois=0.05) and IDR (0.05) cutoff, followed by differential binding analysis to the CNTL αFOXO1 (edgeR_4.0.16, *p*=0.05) with the DiffBind package (https://git.bioconductor.org/packages/DiffBind, version 3.12.0). Consensus H3K27ac peaks were identified as shared peaks in the two replicates that intersected using BEDTools with MACS2 *p*-value (Poisson test, ppois=0.05)^81^. H3K27ac differential binding analysis was completed with the DiffBind package and DESeq2 *p*-value cutoff of 0.05. Motif identification and analysis were completed with HOMER^82^, peak overlap was determined with BEDTools, and heat maps and profile plots were generated with deepTools^83^. FASTQ files for adult zebrafish brain and muscle H3K27ac ChIP-seq were obtained from Yang and Luan et al., 2020 (GSE134055)^62^ and analyzed with the ENCODE ChIP-seq pipeline (https://github.com/ENCODE-DCC/chip-seq-pipeline2) and the previously mentioned computational tools.

### Assay for Transposase-Accessible Chromatin (ATAC-seq)

Embryos were injected with PAX3::FOXO1-2A-sfGFP or pCS2+MCS-P2A-sfGFP mRNA and dissociated into single cells as described above. Omni-ATAC-seq was adapted from Corces et al., 2017^84^ and completed in duplicates from different injection days. 50,000 cells were taken and lysed in 50 μL of 1M Tris-HCl pH 7.4, 10 mM NaCl, 3 mM MgCl_2_ with 0.1% NP-40, 0.1% Tween-20, and 0.01% Digitonin. Cells were incubated on ice for 3 minutes before adding 1 mL of 1M Tris-HCl pH 7.4, 10 mM NaCl, 3 mM MgCl_2_, and 0.1% Tween-20. Samples were centrifuged at 4°C, 500g for 10 min, then the supernatant was removed. 50 μL of transposition mix (25 μL 2X TD buffer (Diagenode, C01019043), 2.5 μL Tn5 (Diagenode, C01070012), 0.5 μL 10% Tween-20, 0.5 μL 1% Digitonin, 16.5 μL dPBS, 5 μL H_2_O) was added and the reaction was incubated at 37°C for 30 minutes in a ThermoMixer at 1000 rpm. DNA was purified with the QIAGEN MinElute PCR Purification kit and eluted in 22 μL EB. DNA libraries were amplified by qRT-PCR until the reaction plateaued. The reaction consisted of 25 μL Phusion High-Fidelity PCR Master Mix (ThermoFisher Scientific, F531L), 5 μL 25 μM unique indexing primer sets from Buenrostro et al., 2013^85^, 0.3 μL 100X SYBR Green I, and 20 μL DNA. The PCR reaction was purified by 0.4X/1.2X double-sided AMPure bead size selection and eluted in 20 μL EB. An additional 1.2X AMPure selection was done to remove excess adapter dimer, which was evaluated by running 1 μL of DNA on an E-gel. Sequencing on the NovaSeq SP with 2×150 bp and additional QC was completed by IGM.

ATAC-seq FASTQ files were analyzed through the ENCODE ATAC-seq pipeline (https://github.com/ENCODE-DCC/atac-seq-pipeline). Bigwigs were generated by deepTools and normalized by Bins Per Million mapped reads (BPM, --binSize 1). Differential accessibility was determined with DiffBind with motif analysis and heat map generation completed as done for ChIP-seq. Publicly available ATAC-seq datasets from Pálfy and Schulze et al., 2020 (GSE130944)^40^ and Liu and Wang et al., 2018 (GSE101779)^39^ were downloaded and analyzed through the ENCODE ATAC-seq pipeline. PCA was completed with DiffBind and accessibility overlap with PAX3::FOXO1 binding sites was determined by BEDTools. ATAC-seq and H3K27ac ChIP-seq were integrated into the activity-by-contact model for enhancer-promoter interactions^42^.

### RNA-sequencing (RNA-seq)

A minimum of 30 embryos injected with either PAX3::FOXO1-2A-sfGFP or pCS2+MCS-P2A-sfGFP mRNA were collected per condition. Embryos were dechorionated with pronase, snap-frozen with dry ice, and stored at −80°C until RNA extraction. This was repeated across two injection days for quadruplicates. RNA isolation and library generation was completed as described in Kent et al., 2023^44^. RNA-seq analysis was performed with an in-house pipeline. In brief, FASTQC (http://www.bioinformatics.babraham.ac.uk/projects/fastqc) and MultiQC^86^ assessed quality, sequences were trimmed for quality and adapter content with TrimGalore (https://github.com/FelixKrueger/TrimGalore). Reads were aligned to danRer11 with STAR^87^ and assigned to genomic features with featureCount^88^. Differential gene expression was determined by DESeq2^89^ and SARTools^90^. Pathway analysis was completed using clusterProfiler^91^, enrichr^92^, the MSigDB database^93^, and DAVID (https://david.ncifcrf.gov/)^94^. A volcano plot was generated by enhancedVolcano (https://github.com/kevinblighe/EnhancedVolcano). RNA-seq FASTQ files were downloaded from White et al., 2017 (PRJEB7244)^43^ and run through the same in-house pipeline. PCA was completed with DiffBind, and pathway analysis with clusterProfiler.

### Statistical analysis and Data Visualization

GraphPad Prism 10 and R (version 4.3.0) were used for all statistical analysis, further data visualization, and gene overlap analysis. Various plots were generated by ggplot2^95^. Sample sizes, the number of times the experiment was repeated, and statistical tests are included in the figure legends.

## Data Availability

PAX3::FOXO1 ChIP-seq, ATAC-seq, H3K27ac ChIP-seq, and RNA-seq datasets will be deposited in the NCBI Gene Expression Omnibus (GEO, http://www.ncbi.nlm.nih.gov/geo/). Previously published zebrafish ATAC-seq datasets from embryonic zebrafish development were obtained from <u>GSE130944</u>^40^ and <u>GSE101779</u>^39^. Embryonic zebrafish RNA-seq datasets were obtained from the project <u>PRJEB12296</u>^43^ in the European Nucleotide Archive (ENA, https://www.ebi.ac.uk/ena). ATAC-seq and H3K27ac ChIP-seq in adult zebrafish muscle and brain tissues were obtained from GSE134055^62^.

## Code Availability

This paper reports a new PerCell ChIP-seq analysis pipeline, which was utilized for PAX3::FOXO1 and H3K27ac ChIP-seq experiments. Code for this pipeline can be found at https://github.com/lextallan/PerCell. Information about this pipeline can be found in Tallan et al., 2024. This paper also utilized an in-house RNA-seq pipeline, which was described in the methods section. The code used for this paper will be made available on Github (https://github.com/Kendall-Cancer-Lab/PAX3-FOXO1_invivo_6hpf). Any additional information needed to reanalyze this data is available from the corresponding authors.

## Acknowledgements

We thank the Nationwide Children’s Hospital (NCH) Animal Resources Core for their exceptional zebrafish husbandry, especially the Zebrafish Facility team members Dr. Laurie Goodchild, Dr. Carmen Arsuaga, Dr. Lindsey Ferguson, Logan Fehrenbach, Alex Kramer, and Logan Bern. Additional support was provided by the NCH Flow Cytometry Core, the NCH Institute for Genomic Medicine, the NCH Genomics Services Laboratory, and the NCH High-Performance Computing group for assistance maintaining and using the NCH cluster. We also thank Dr. Matthew Kent for assistance with cloning and Dr. Benjamin Sunkel for ChIP-seq training. We are grateful to all members of the Stanton and Kendall groups for helpful discussions.

G.C.K. is grateful for support from an NIH/NCI R01 CA272872 grant, an Alex’s Lemonade Stand Foundation “A” Award, a V Foundation for Cancer Research V Scholar Award, a CancerFree Kids New Idea Award, and Startup Funds from The Abigail Wexner Research Institute at Nationwide Children’s Hospital. B.Z.S. is grateful to the American Cancer Society (RSG-23-1021178-01-DMC), St. Baldrick’s Foundation (Career Development Award), National Institutes of Health (R01GM144601, 1R01HL166520 - 01A1), and intramural funding from Nationwide Children’s Hospital for supporting this work. G.C.K and B.Z.S. were supported by a Nationwide Children’s Hospital Seed Fund from the Center for Childhood Cancer Research. J.K. is supported by a T32 CA269052 Training Program in Basic and Translational Pediatric Oncology Research predoctoral fellowship. The Institute for Genomic Medicine is funded by the Nationwide Foundation Pediatric Innovation Fund and the Ohio State University Comprehensive Cancer Center grant P30 CA016058. The funders had no role in study design, data collection and analysis, decision to publish, or preparation of the manuscript. Further, the content is solely the responsibility of the authors and does not necessarily represent the official views of the National Institutes of Health.

## Author Information

### Authors and Affiliations

**Molecular, Cellular, and Developmental Biology Ph.D. Program, The Ohio State University, Columbus, OH, USA**

Jack Kucinski, Alexi Tallan, Benjamin Z. Stanton, and Genevieve C. Kendall

**Center for Childhood Cancer Research, The Abigail Wexner Research Institute, Nationwide Children’s Hospital, Columbus, OH, USA**

Jack Kucinski, Alexi Tallan, Cenny Taslim, Meng Wang, Matthew V. Cannon, Katherine M. Silvius, Benjamin Z. Stanton, and Genevieve C. Kendall

**Department of Pediatrics, The Ohio State University College of Medicine, Columbus, OH, USA**

Benjamin Z. Stanton and Genevieve C. Kendall

**Department of Biological Chemistry and Pharmacology, The Ohio State University College of Medicine, Columbus, OH, USA**

Benjamin Z. Stanton

### Contributions

J.K., B.Z.S., and G.C.K. conceptualized this study. J.K. led investigations/experimentation with K.M.S. providing validation studies. Zebrafish methodology was developed by J.K. and PerCell ChIP-seq pipeline methodology/software was developed by A.T. and M.W. Formal analysis was done by J.K., A.T., C.T. M.W., and M.V.C. Data visualization was completed by J.K., A.T., and C.T. Supervision and funding acquisition was done by B.Z.S. and G.C.K. The initial manuscript and figures were drafted by J.K., A.T., K.M.S., B.Z.S., and G.C.K. All authors reviewed and edited the final manuscript.

### Corresponding authors

Correspondence should be addressed to Benjamin Stanton or Genevieve Kendall.

## Ethics declarations

All authors declare no competing interests.

---

**Extended Data Fig. 1:**
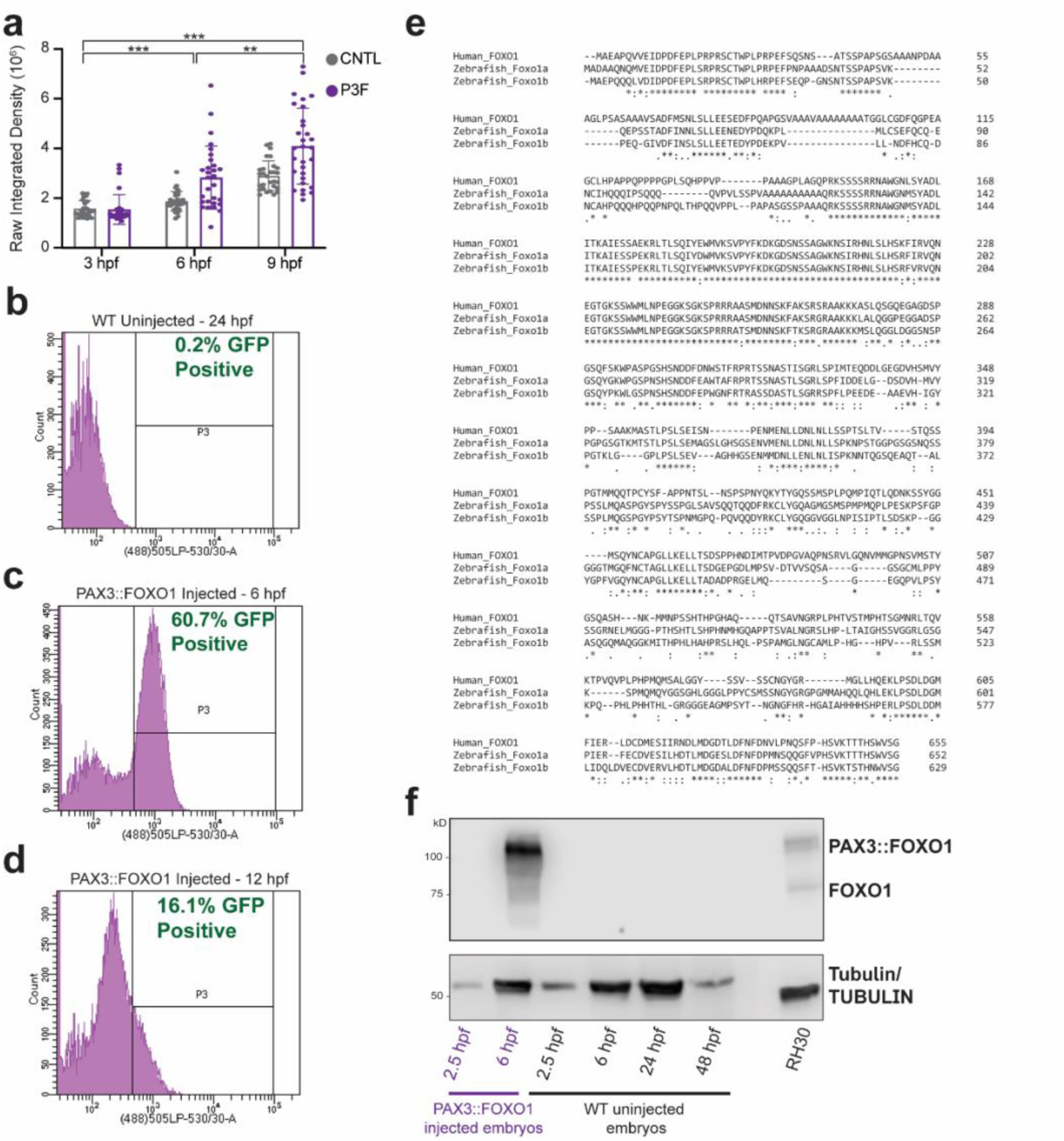
Characterization of PAX3::FOXO1 expression kinetics shows drop-off at later time points. (a) Quantification of raw integrated density of GFP fluorescence. Each point represents an individual embryo, which were collected across three different injection days. Control-injected (CNTL) embryos: n=29 per time point, PAX3::FOXO1-injected (P3F) embryos: n=30 at 3 and 9 developmental hours post-fertilization (hpf), n=29 at 6 hpf. A Brown-Forythe and Welch ANOVA test with a Benjamini, Krieger, and Yekutieli correction for multiple comparisons was used to determine statistical significance. Asterisks represents statistics between P3F embryos. CNTL embryos across time-points had p≤0.005. Error bars indicate standard deviation. (b) Flow cytometry for GFP of 24 hpf wildtype (WT) uninjected zebrafish embryo as negative control. (c, d) Flow cytometry for GFP of zebrafish embryos injected with 50 ng/μL of PAX3::FOXO1 and 50 ng/uL pCS2-tagRFPT.zf1 at 6 and 12 hpf, respectively. (e) Protein sequence BLAST alignment between human FOXO1 and zebrafish orthologs Foxo1a and Foxo1b. (f) Western blot with protein lysate from 15 embryos per lane for each condition and 10 μg of protein lysate from RH30 cells, a PAX3::FOXO1-positive patient-derived cell line. Endogenous FOXO1 is detected in the patient-derived sample but not in zebrafish embryos. WT – Wildtype. *** denotes p<0.001 and ** denotes 0.001<p<0.01.

**Extended Data Fig. 2:**
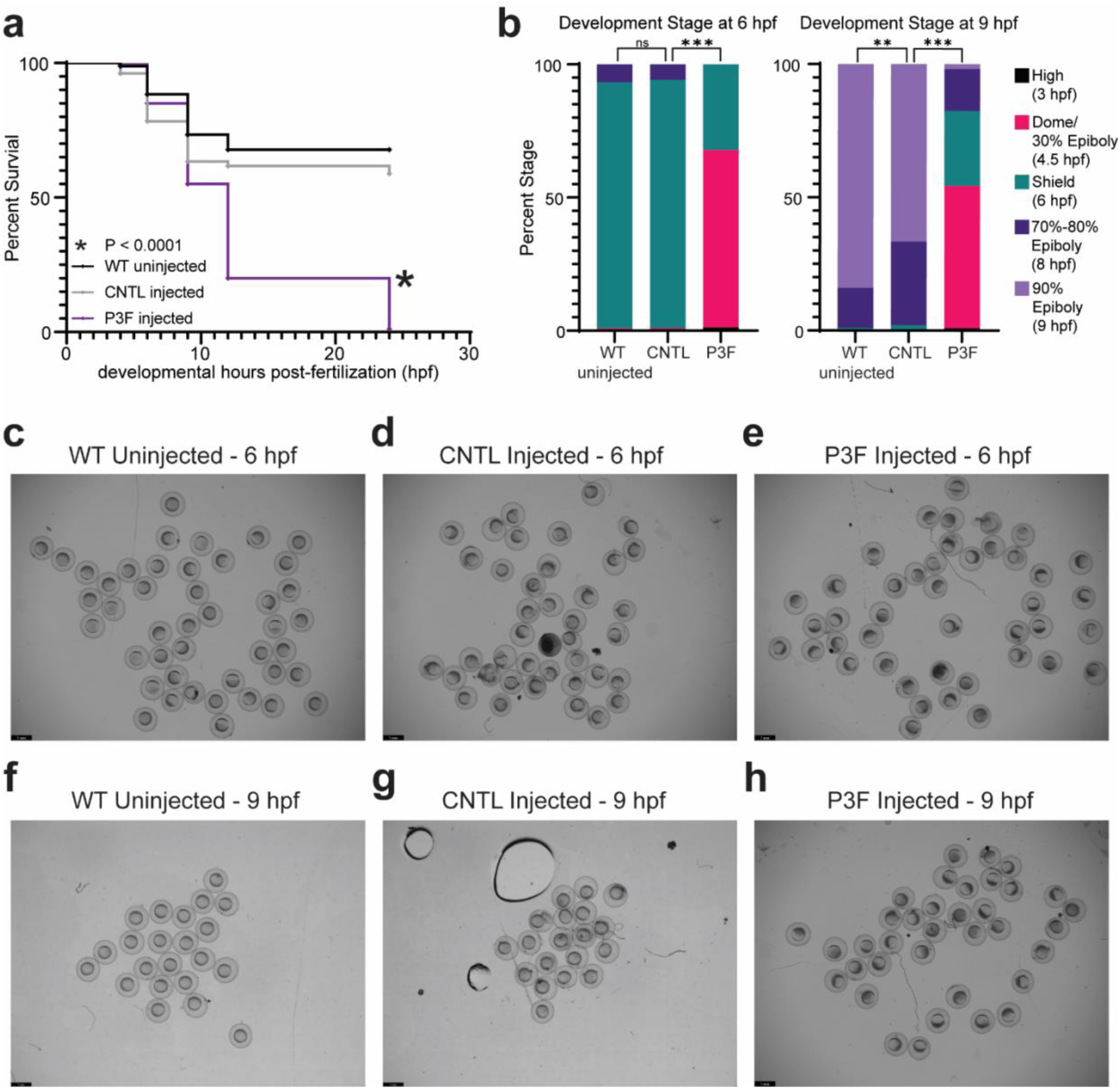
PAX3::FOXO1 causes arrested development in zebrafish embryos. (a) Kaplan-Meier survival curve with log-rank (Mantel-Cox) test with 180 embryos per condition, wildtype uninjected (WT), control-injected (CNTL), or PAX3::FOXO1-injected (P3F) embryos are included across two injection days. (b) Embryos were visually evaluated for developmental stage at certain time points according to phenotypic descriptions in Kimmel et al., 1995^34^. Embryos were scored and data combined across three injection days for a total of WT uninjected 6 hours-post-fertilization (hpf): n=119, WT uninjected 9 hpf: n=93, CNTL 6 hpf: n=104, CNTL 9 hpf: n=160, P3F 6 hpf: n=84, P3F 9 hpf: n=75. A Fisher’s exact test with a Bonferroni correction for multiple comparisons for the number of embryos at the expected developmental stage was used to determine statistical significance. (c-h) Representative field of view images for WT, CNTL, or P3F embryos at 6 and 9 hpf. Scale bar, 1 mm. *** denotes p<0.001 and ** denotes 0.001<p<0.01.

**Extended Data Fig. 3:**
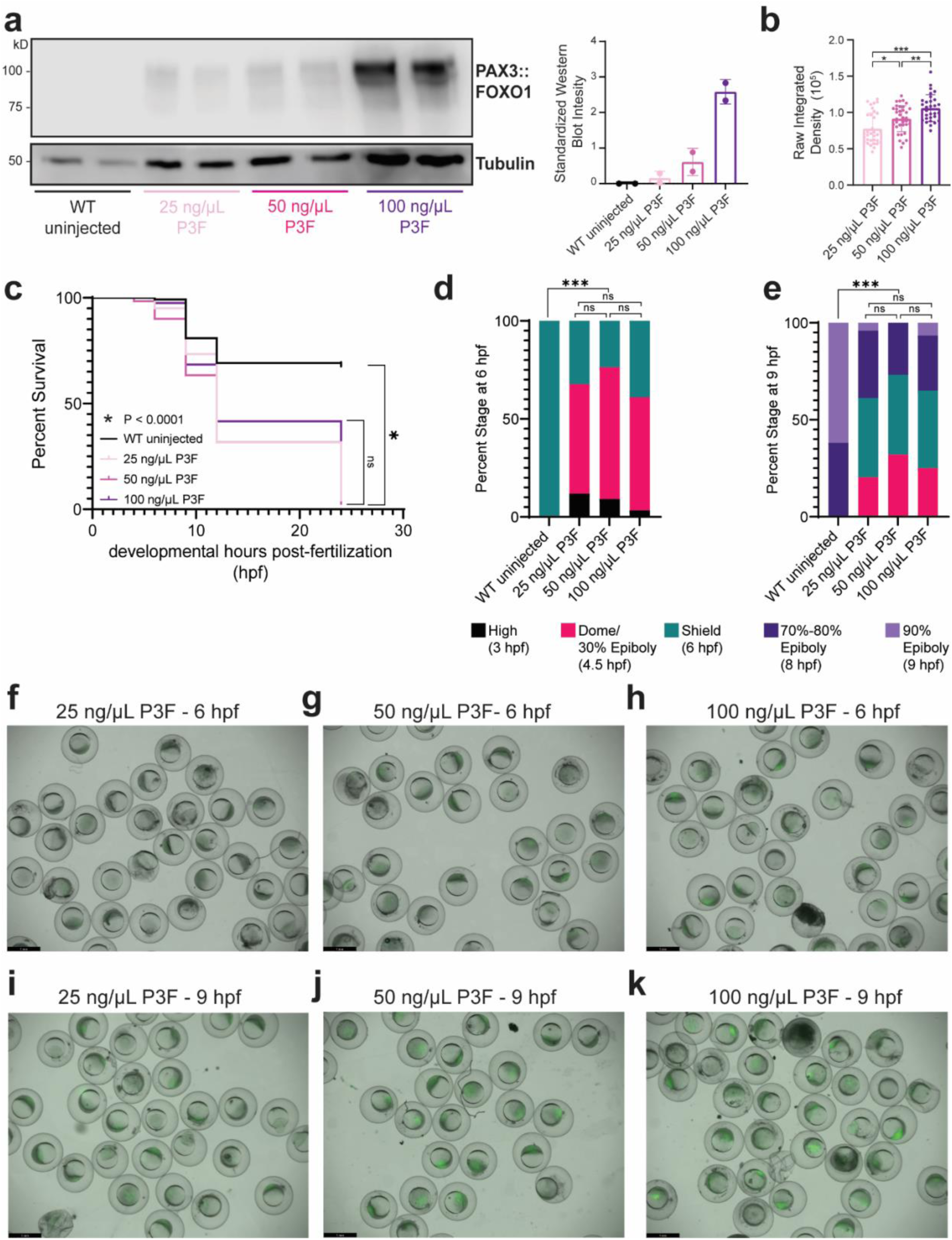
PAX3::FOXO1 injection levels do not alter broad phenotypic consequences. (a) Representative western blot with protein lysate from 12 embryos per lane for each injection concentration of PAX3::FOXO1 (P3F). Normalized quantification for the representative western blot of PAX3::FOXO1/αTUBULIN signal is on the right, n=2 duplicates (presented on left) to generate the mean. The titration western blot was repeated three times from one injection day. Error bars indicate standard deviation. (b) GFP quantification by ImageJ of PAX3::FOXO1-injected (P3F) embryos at various injection concentrations. Each point represents an individual embryo, which were measured across two different injection days. 25 ng/μL P3F: n=33, 50 ng/μL P3F: n=37, 100 ng/μL P3F: n=30. A one-way ANOVA with a Tukey’s test for multiple comparisons was used to determine statistical significance. Error bars indicate standard deviation. (c) Kaplan-Meier survival curve with log-rank (Mantel-Cox) test and Bonferroni correction for multiple comparisons with 120 embryos per condition with wildtype uninjected (WT) and titrating concentrations of 25, 50, or 100 ng/μL of PAX3::FOXO1 mRNA injection (P3F) across two injection days. (d, e) Embryos were visually evaluated for developmental stage at certain time points according to phenotypic descriptions in Kimmel et al., 1995^34^. Embryos were scored and data combined across three injection days for a total of WT 6 hpf: n=81, 25 ng/μL P3F 6 hpf: n=59, 50 ng/μL P3F 9 hpf: n=55, 100 ng/μL P3F 6 hpf: n=59, WT 9 hpf: n=84, 25 ng/μL P3F 9 hpf: n=49, 50 ng/μL P3F 9 hpf: n=56, 100 ng/μL P3F 9 hpf: n=60. (f-k) Representative field of view images of 25 ng/μL P3F, 50 ng/μL P3F, 100 ng/μL P3F. Scale bar, 1 mm. *** denotes p<0.001, ** denotes 0.001<p<0.01, and * denotes 0.01<p<0.05 unless otherwise noted.

**Extended Data Fig. 4:**
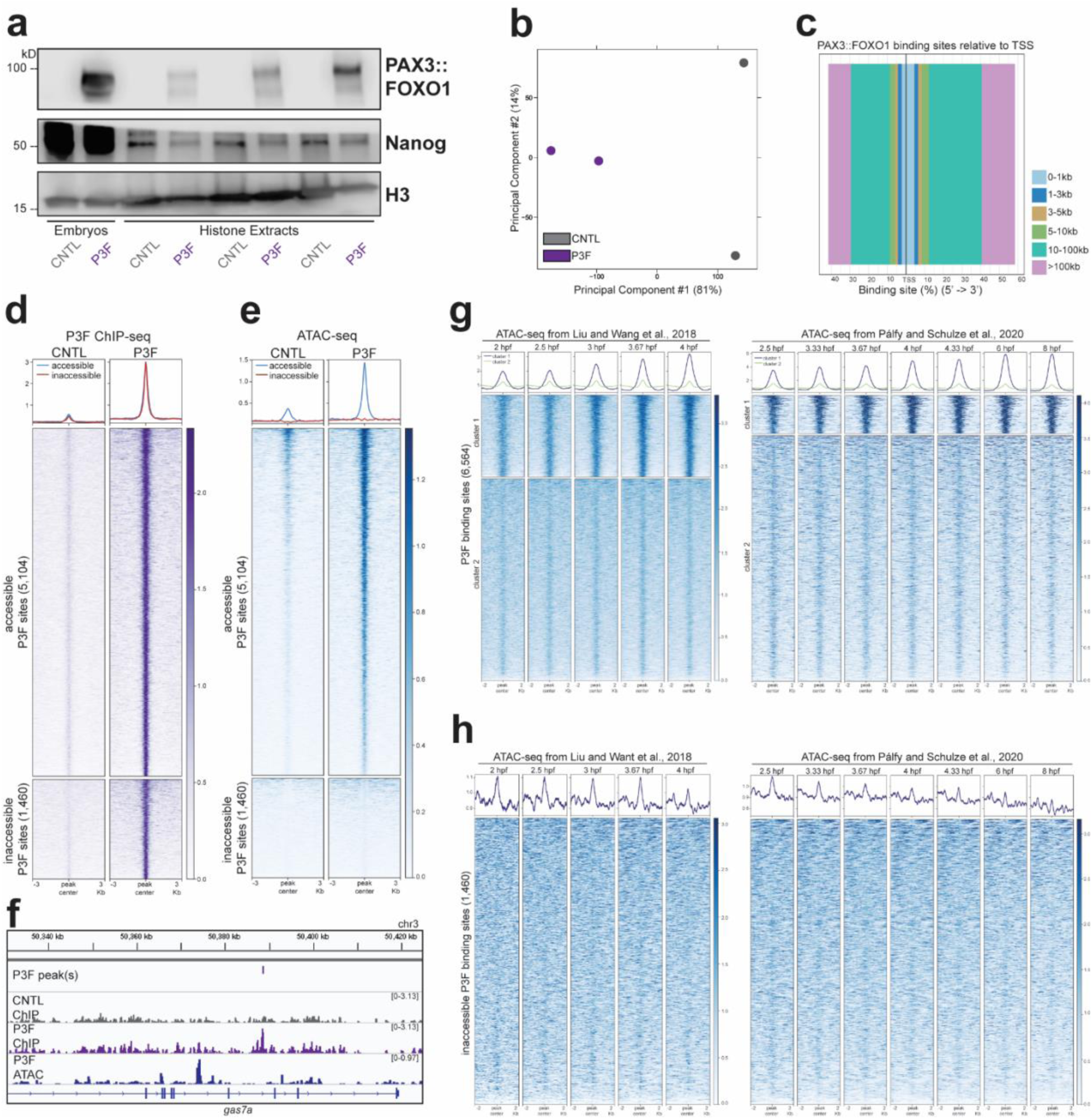
PAX3::FOXO1 binding occurs at sites of closed chromatin. (a) Western blot on whole embryos (far left) and histone extractions for protein DNA interaction by blotting for PAX3::FOXO1, Nanog (pioneer transcription factor), and histone H3 in control-injected (CNTL) and PAX3::FOXO1-injected (P3F) embryos. Samples were collected across two injection days. (b) Principal component analysis (PCA) on consensus peaks between replicates of αFOXO1 ChIP-seq in CNTL and P3F embryos. (c) Distance of PAX3::FOXO1 binding sites to nearest zebrafish transcriptional start site (TSS) by genomic distance. (d, e) Average PAX3::FOXO1 ChIP-seq and ATAC-seq signal between replicates at accessible sites that overlap with an ATAC-seq peak (top) and inaccessible sites without an ATAC-seq peak (bottom), respectively. (f) Representative Integrative Genome Viewer (IGV) tracks of intronic PAX3::FOXO1 binding in *gas7a* that lacks accessibility. (g) Merged ATAC-seq signal across replicates at PAX3::FOXO1 binding sites during zebrafish embryonic development according to hours post-fertilization (hpf) from Liu and Wang et al., 2018^39^ and Pálfy and Schultze et al., 2020^40^ with k-means clustering (k=2). (h) Merged ATAC-seq signal across replicates at inaccessible PAX3::FOXO1 binding sites from Fig. 2c during zebrafish embryonic development according to hpf from Liu and Wang et al., 2018^39^ and Pálfy and Schultze et al., 2020^40^.

**Extended Data Fig. 5:**
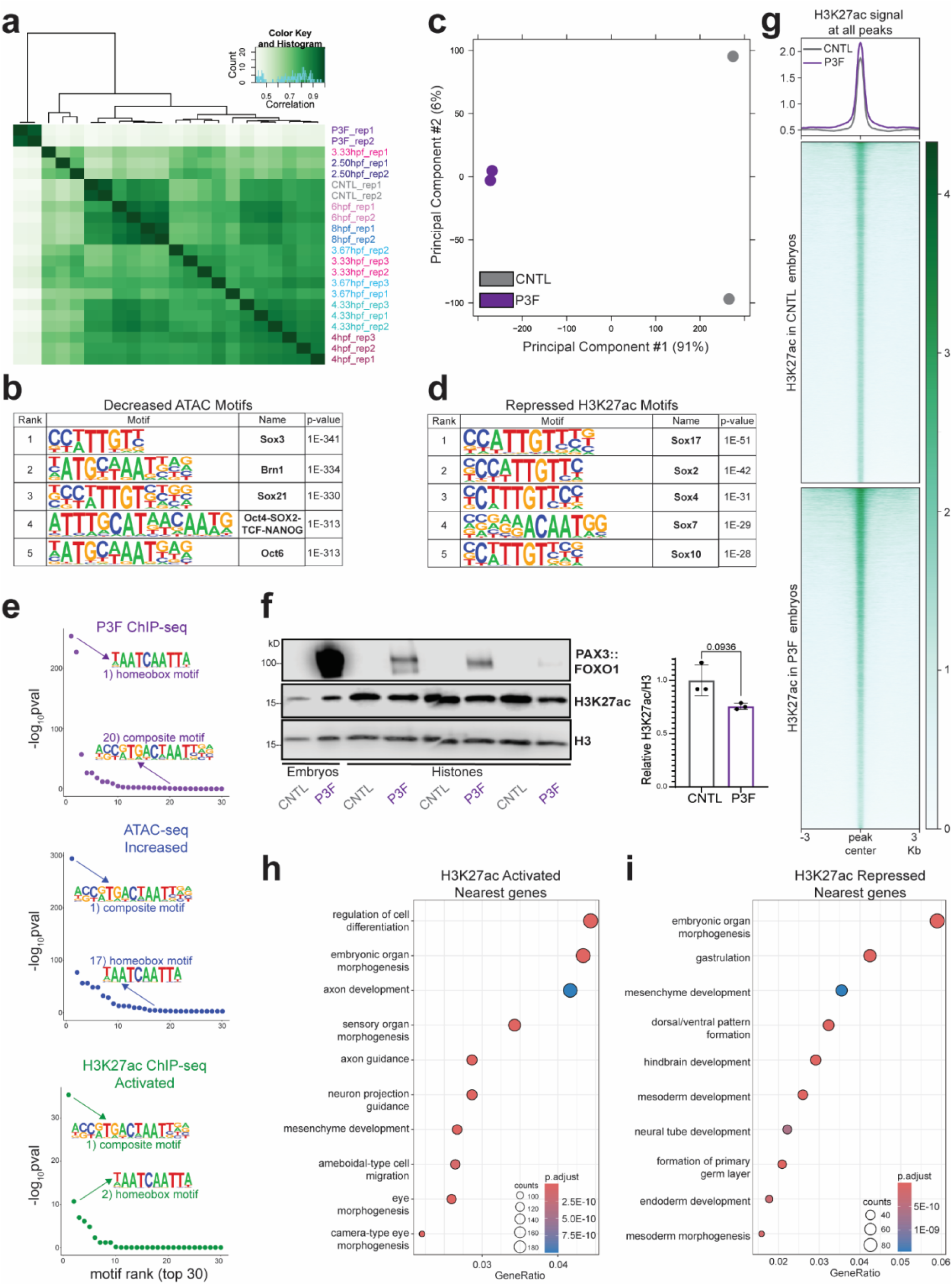
PAX3::FOXO1 redistributes active chromatin features. (a) Hierarchal clustering of ATAC-seq principal component analysis (PCA) shown in Fig. 4a. (b) Top five known HOMER motifs at ATAC-seq peaks with decreased accessibility in PAX3::FOXO1-injected (P3F) embryos. (c) PCA on consensus H3K27ac ChIP-seq replicate peaks in control-injected (CNTL) and P3F embryos. (d) Top five known HOMER motifs at sites with lower H3K27ac ChIP-seq signal in P3F embryos. (e) Relative enrichment of composite and homeobox motifs as described in Fig. 3g at PAX3::FOXO1 binding sites (purple), sites with increased ATAC-seq signal (blue), and higher H3K27ac ChIP-seq signal (green). (f) Western blot on whole embryos (far left) and histone extractions for PAX3::FOXO1, active histone mark H3K27ac, and histone H3. PAX3::FOXO1 and H3 blots are the same as in Figure 2b, as these experiments were completed simultaneously. This experiment was replicated with six samples per condition across three injection days. Normalized quantification for the representative western blot of H3K27ac/H3 signal is compared to the average signal ratio in CNTL. An unpaired Welch’s t-test was used to determine statistical significance. (g) H3K27ac ChIP-seq signal across all called peaks in each replicate and condition. (h-i) Top 10 enriched gene ontology biological pathways on genes nearest to sites with higher H3K27ac and lower H3K27ac in P3F embryos, respectively.

**Extended Data Fig. 6:**
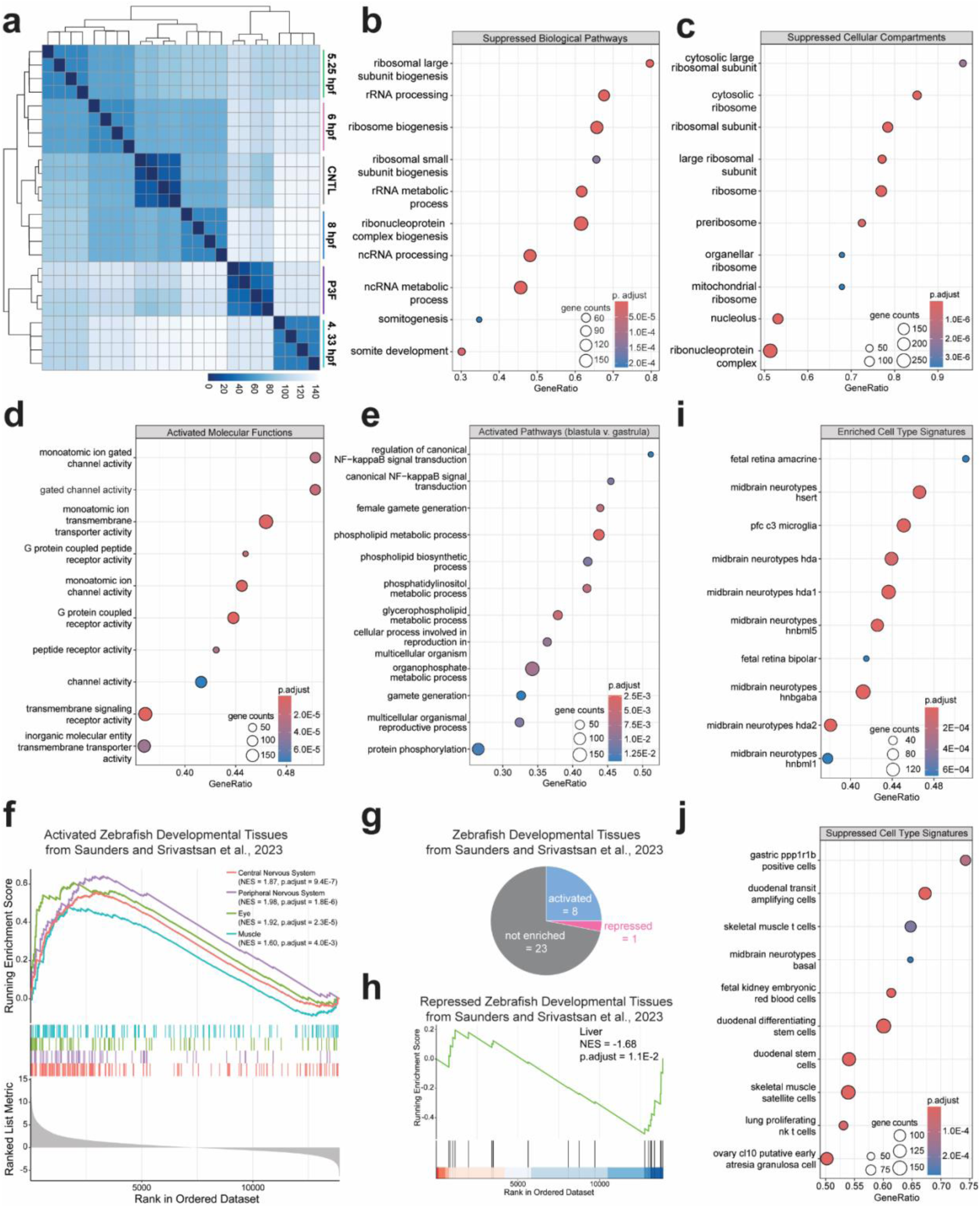
Transcriptional upregulation of neural cell state pathways and downregulation of ribosomal and developmental signatures in PAX3::FOXO1-injected embryos. (a) Hierarchal clustering of RNA-seq PCA shown in Fig. 5a. (b) Gene set enrichment analysis (GSEA) for downregulated gene ontology biological pathways in PAX3::FOXO1-injected (P3F) embryos. (c) GSEA for downregulated gene ontology cellular compartments in P3F embryos. (d) GSEA for upregulated gene ontology molecular functions in P3F embryos. (e) GSEA for upregulated gene ontology biological pathways in blastula stage zebrafish embryos at 4.33 hours post-fertilization (hpf) versus gastrula stage zebrafish embryos at 6 hpf. (f) GSEA to markers of the top 4 most enriched developing zebrafish tissues identified from scRNA-seq analysis from Saunders and Srivatsan et al., 2023^51^. (g) Overview of GSEA analysis for markers of all developing zebrafish tissues identified from scRNA-seq analysis from Saunders and Srivatsan et al., 2023^51^. (h) GSEA to markers of developing zebrafish liver tissue from Saunders and Srivatsan et al., 2023^51^. (i-j) GSEA for cell type signatures from the Molecular Signatures Database (MSigDB) for upregulated and downregulated signatures in P3F embryos, respectively.

**Extended Data Fig. 7:**
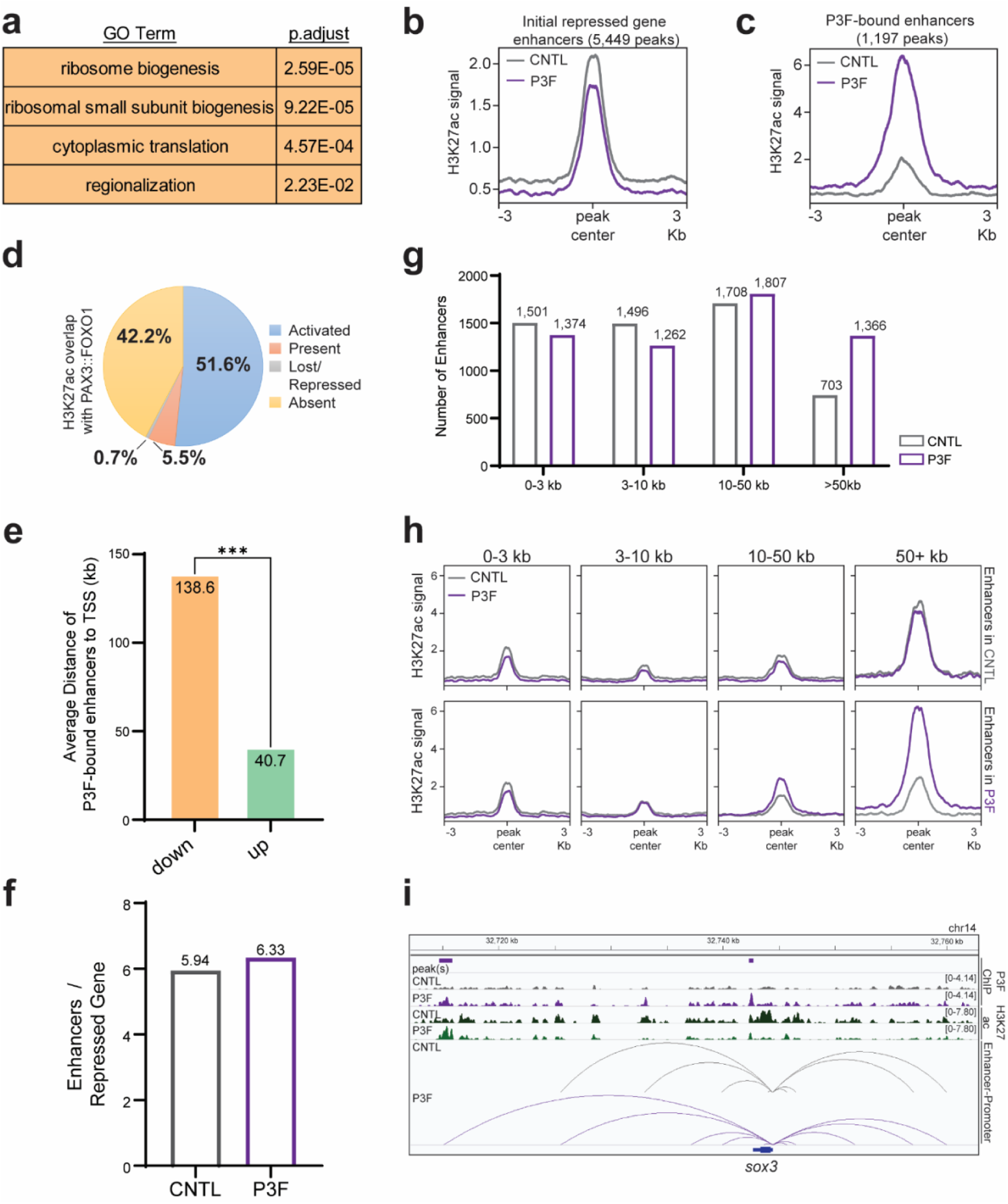
PAX3::FOXO1 weakens nearby enhancers to repress genes. (a) The top 5 enriched gene ontology biological pathways with DAVID analysis for genes directly-downregulated by PAX3::FOXO1, n=917. (b) Average H3K27ac ChIP-seq signal between replicates at sites that regulated genes directly downregulated by PAX3::FOXO1 in control-injected (CNTL) embryos according to the activity-by-contact model. (c) Average H3K27ac ChIP-seq signal between replicates at PAX3::FOXO1-bound sites that directly downregulate genes. (d) Overlap of H3K27ac peak characterization at PAX3::FOXO1 binding sites. (e) Average genomic distance of PAX3::FOXO1-bound H3K27ac peaks to downregulated genes (n=1,197 interactions) and upregulated genes (n=10,910 interactions). A Kolmogorov-Smirnov test was used to determine statistical significance. (f) Average number of enhancers that regulate genes directly downregulated by PAX3::FOXO1 in CNTL and PAX3::FOXO1-injected (P3F) embryos. (g) Number of enhancers regulating genes directly downregulated by PAX3::FOXO1 in CNTL and P3F embryos binned by genomic distance to transcriptional start site. (h) H3K27ac ChIP-seq signal for enhancers binned in Extended Data Fig. 7g. (i) Representative IGV tracks near *sox3*. Top tracks are PAX3::FOXO1 ChIP-seq (purple) and middle tracks are H3K27ac ChIP-seq (green). Gray lines indicate predicted enhancer-promoter in CNTL embryos and purple lines indicate predicted enhancer-promoter interactions in P3F embryos for *sox3*. *** denotes p<0.001.

**Extended Data Fig. 8:**
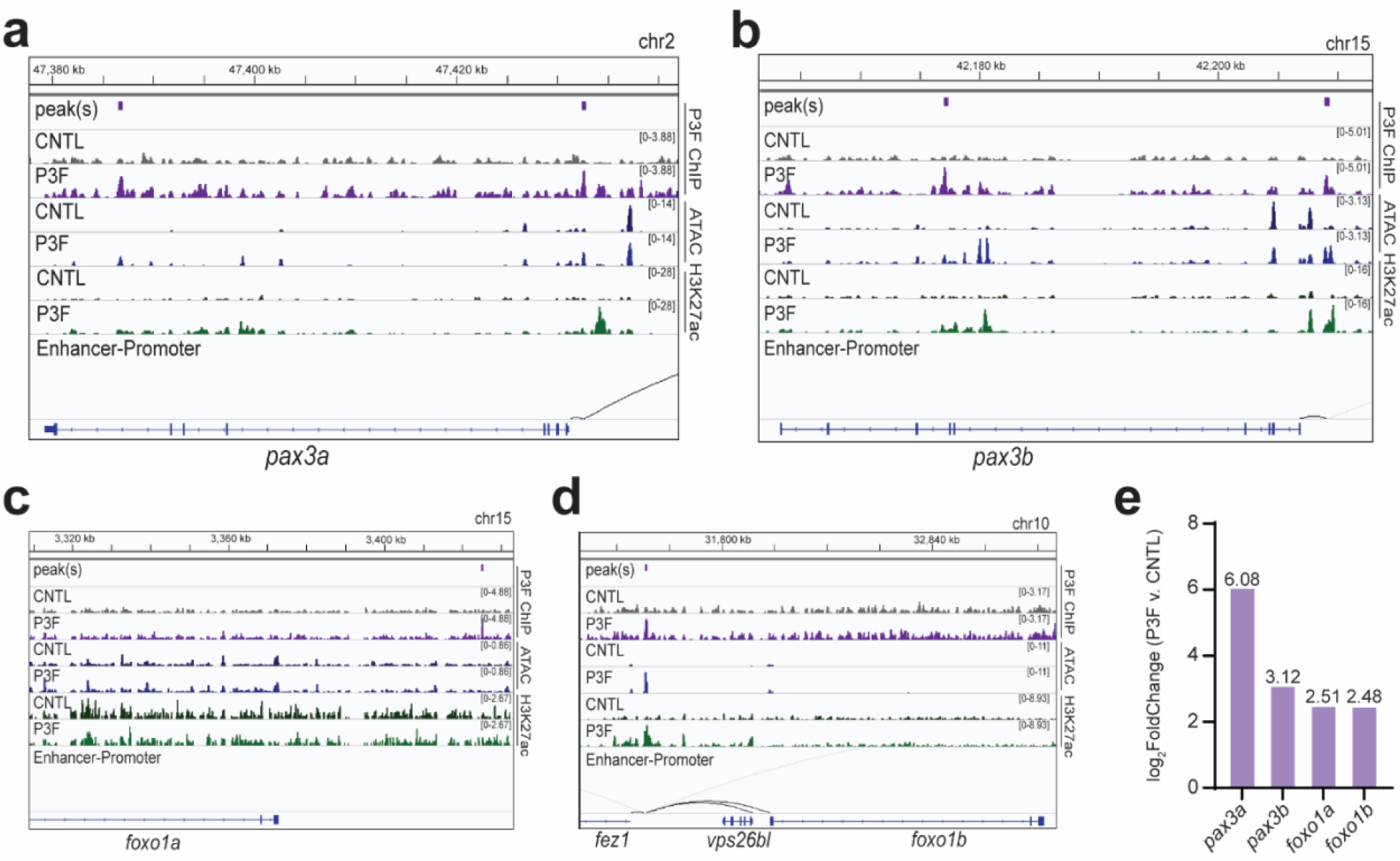
PAX3::FOXO1 activates endogenous fusion partners. (a-d) IGV tracks showing PAX3::FOXO1 binding and regulation of the endogenous fusion partners in zebrafish of *pax3a*, *pax3b*, *foxo1*, and *foxo1b*, respectively. Top tracks are PAX3::FOXO1 ChIP-seq (purple), middle tracks are ATAC-seq (blue), and lower tracks are H3K27ac ChIP-seq (green). Black lines indicate predicted enhancer-promoter interactions in P3F embryos. (e) Average RNA-seq log_2_ fold change of endogenous fusion partner expression in PAX3::FOXO1-injected (P3F) versus control-injected (CNTL) embryos. All fusion partners were differentially expressed from RNA-seq.

**Extended Data Fig. 9:**
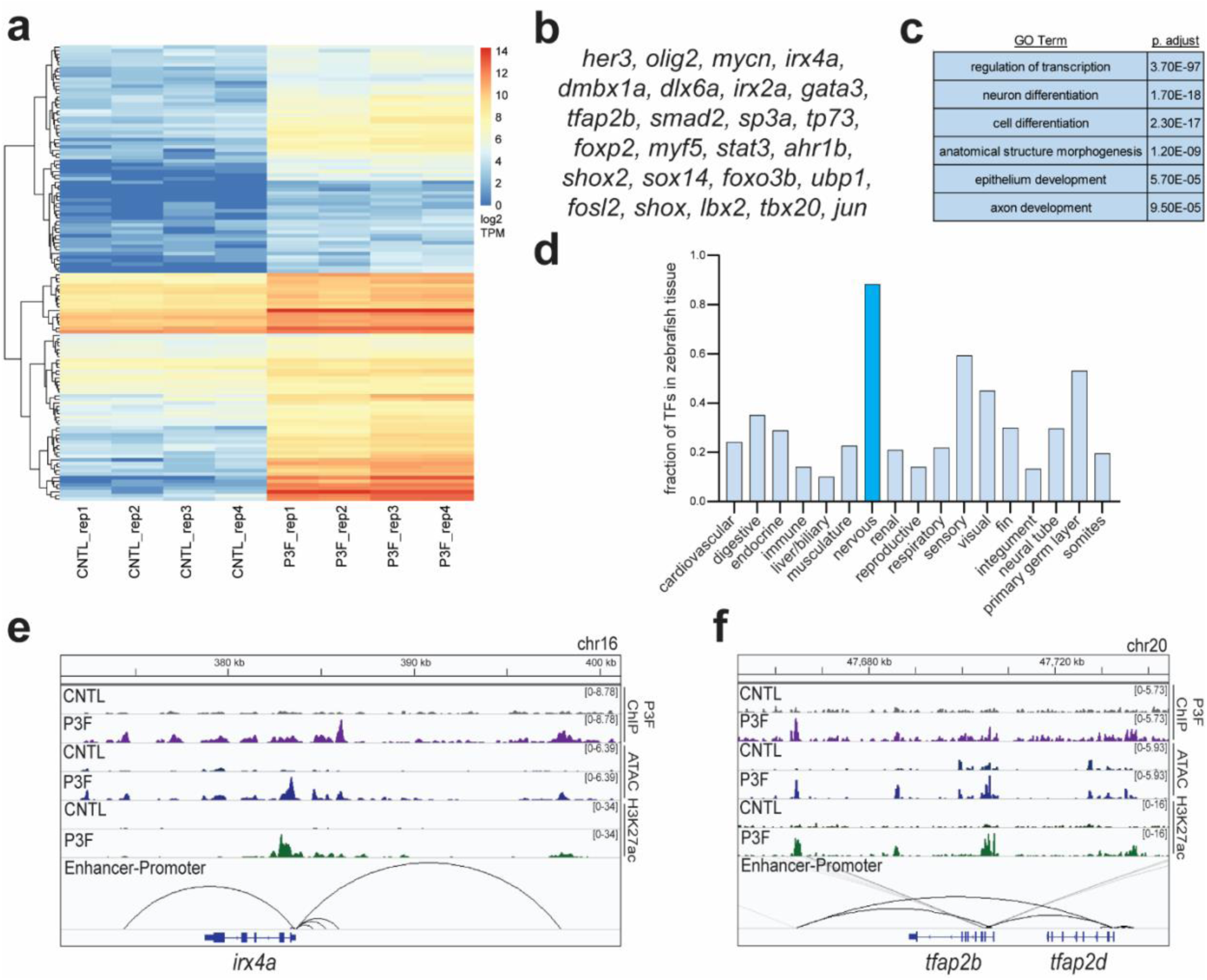
PAX3::FOXO1 activates neural transcription factors. (a) Expression of 130 transcription factors by log_2_(TPM+1) that are directly-upregulated by PAX3::FOXO1 in each RNA-seq replicate of control-injected (CNTL) and PAX3::FOXO1-injected (P3F) embryos. (b) Top 25 most highly expressed directly-upregulated transcription factors by PAX3::FOXO1. (c) The top 5 enriched gene ontology biological pathways with DAVID analysis for transcription factors directly-upregulated by PAX3::FOXO1. (d) Confirmed anatomical structure expression of directly-upregulated transcription factors by PAX3::FOXO1 according to the ZFIN (zfin.org) gene expression data. (e, f) Representative IGV tracks of activation of neural zebrafish transcription factors *irx4a* and *tfap2b*, respectively. Top tracks are PAX3::FOXO1 ChIP-seq (purple), middle tracks are ATAC-seq (blue), and lower tracks are H3K27ac ChIP-seq (green). Black lines indicate new predicted enhancer-promoter interactions for *irx4a* or *tfap2b* in P3F embryos.

**Extended Data Fig. 10:**
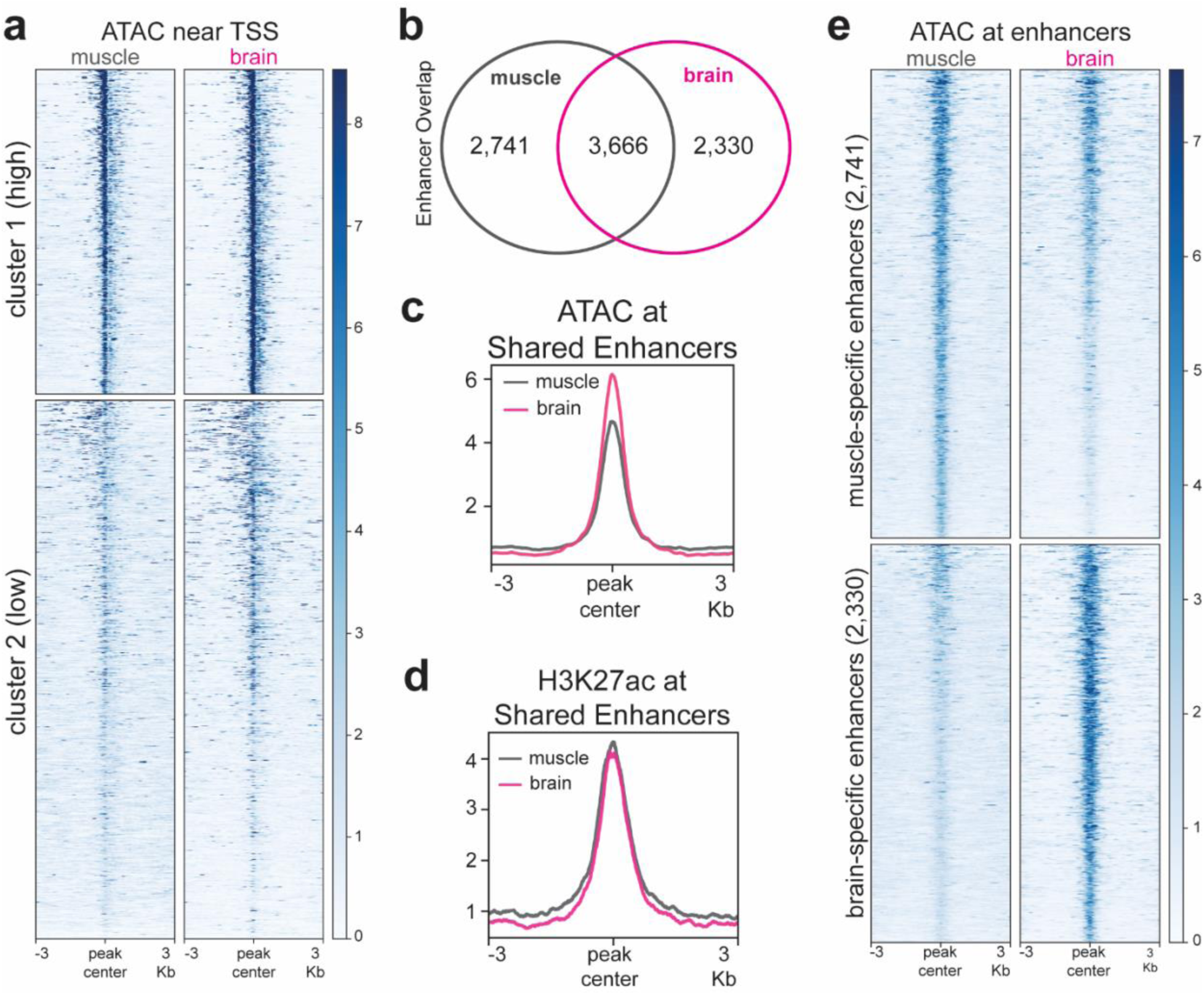
Initial PAX3::FOXO1 targets are more active in brain versus muscle tissue. (a) Heatmap representation of ATAC-seq data presented in Fig. 6g. (b) Overlap of enhancers regulating genes directly-upregulated by PAX3::FOXO1 according to the activity-by-contact model in adult zebrafish muscle and brain tissue. ATAC-seq and H3K27ac ChIP-seq is from Yang and Luan et al., 2020^62^. (c) Average ATAC-seq signal between replicates of shared enhancers of genes directly-upregulated by PAX3::FOXO1. (d) Average ATAC-seq signal between replicates of shared enhancers of genes directly-upregulated by PAX3::FOXO1. (e) Average H3K27ac ChIP-seq signal between replicates of enhancers specific to zebrafish brain or muscle tissue that regulate genes directly-upregulated by PAX3::FOXO1.

**Supplemental Table 1.**
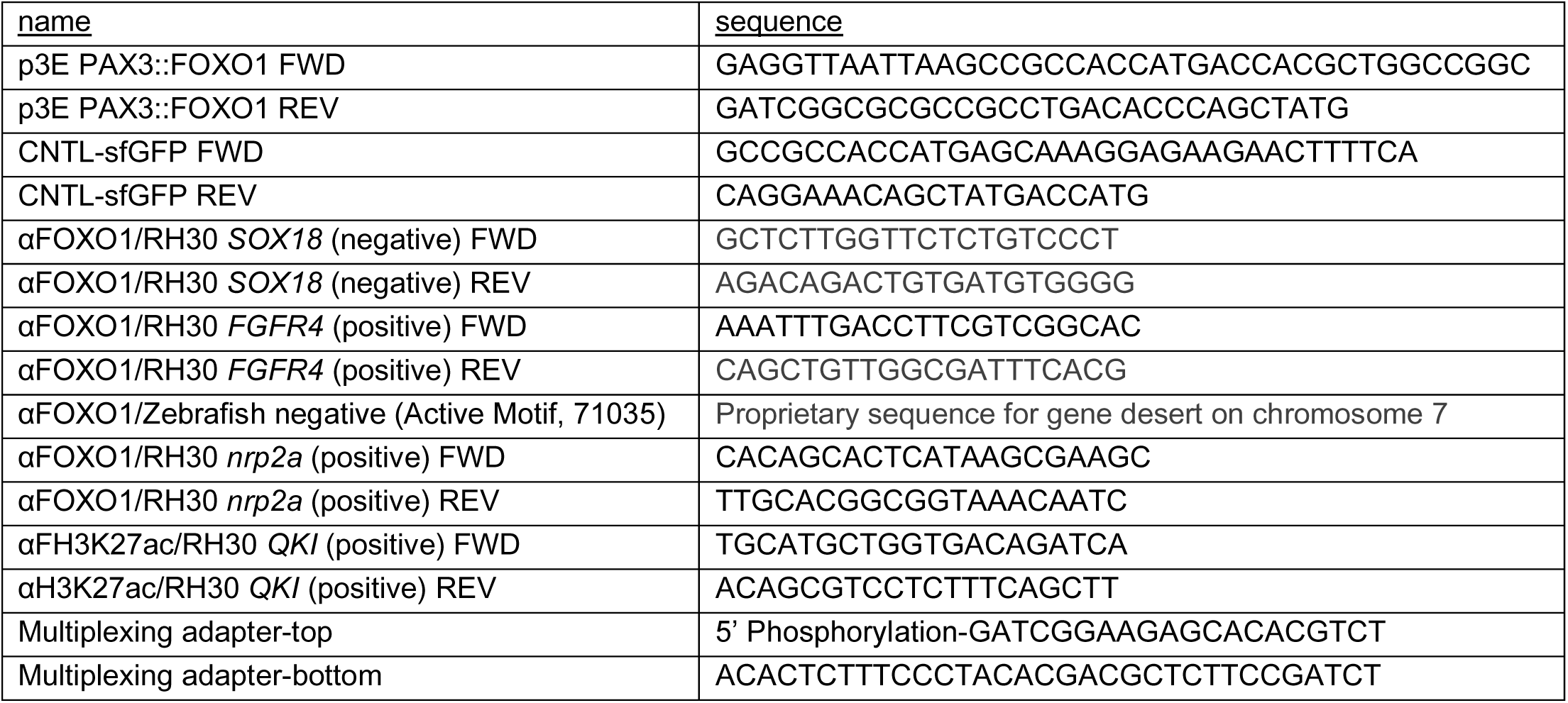
Primers used in this study.

